# A role for liquid-liquid phase separation in ESCRT-mediated nuclear envelope reformation

**DOI:** 10.1101/577460

**Authors:** Alexander von Appen, Dollie LaJoie, Isabel E. Johnson, Mike Trnka, Sarah M. Pick, Alma L. Burlingame, Katharine S. Ullman, Adam Frost

**Affiliations:** Department of Biochemistry and Biophysics, University of California, San Francisco, CA, USA; Department of Oncological Sciences, Huntsman Cancer Institute, University of Utah, Salt Lake City, UT, USA; Department of Pharmaceutical Chemistry, University of California, San Francisco, San Francisco, CA, USA; Faculty of Chemistry and Pharmacy, University of Freiburg, Freiburg, Germany; Chan Zuckerberg Biohub, San Francisco, CA, USA; Quantitative Biosciences Institute, University of California, San Francisco, CA USA

**Author notes:** A.v.A., D.L., and I.E.J. contributed equally to this work.

## Abstract

At mitotic exit, microtubule arrays are dismantled in concert with the reformation of the nuclear envelope. We show how the inner nuclear membrane protein, LEM2, exploits liquid-liquid phase separation to direct microtubule remodeling and nuclear envelope sealing via the Endosomal Sorting Complexes Required for Transport (ESCRT) pathway. LEM2 tethers membrane to chromatin disks through direct binding between its LEM motif and the chromatin-associated barrier-to-autointegration factor (BAF). Concurrently, a low-complexity domain within LEM2 undergoes liquid-liquid phase separation to coat spindle microtubule bundles. Spatially restricted, LEM2’s winged helix (WH) domain activates the ESCRT-II/ESCRT-III hybrid protein, CHMP7. Together LEM2 and CHMP7 copolymerize around microtubule bundles to form a molecular “O-ring” that promotes nuclear compartmentalization and initiates downstream ESCRT factor recruitment. These results demonstrate how multivalent interactions of a transmembrane protein, including those that mediate phase separation, coordinate localized ESCRT polymerization, mitotic spindle disassembly, and membrane fusion. Defects in this pathway compromise spindle disassembly, nuclear integrity, and genome stability.

---

At the onset of “open” mitosis in mammalian cells, the nuclear envelope (NE) dissociates from the chromatin surface to allow for chromosome singularization, which enables the mitotic spindle to segregate chromosomes into emerging daughter cells^1^. In parallel, mitotic phosphorylation events disrupt the affinity of inner nuclear membrane (INM) proteins for the chromatin surface, allowing them to recede, along with NE membranes, into the contiguous endoplasmic reticulum^2^. Upon mitotic exit during anaphase, a program of phosphatases reverses this process: INM proteins within the endoplasmic reticulum membrane regain affinity for DNA or DNA-associated partners to coat each mass of condensed chromosomes, referred to as a chromatin disk, with membranes^3^.

During nuclear envelope reformation, barrier-to-autointegration factor (BAF) cross-bridges anaphase DNA in a meshwork that restricts membrane access to the surface of the disk and guides the formation of a single nucleus with the correct chromosomal composition in a process that lasts approximately 12 minutes^4,5^. As the nascent nuclear envelope engulfs the chromatin disk, a subset of the nuclear membrane proteome transiently segregates into a distinct core region that coincides with microtubules emanating from centrosomes on the outer face of the disk and with central spindle microtubules on the inner face of the disk^4,6–8^. Of the INM proteins that enrich at the core, several belong to the LEM (Lap2-Emerin-MAN1) domain family of proteins, which are characterized by their interactions with the nuclear lamina and, with respect to Lap2 and Emerin, their ability to bind to BAF via a conserved LEM-domain^9^. Recent studies identified another LEM family protein, the two pass transmembrane protein LEM2^10^, as the site-specific adaptor for the ESCRT machinery^11,12^. The ESCRT pathway ultimately coordinates spindle clearance and nuclear envelope fusion for the reforming nucleus^6,13^.

Within the core region of reforming nuclei, spindle disassembly and nuclear envelope (NE) fusion occur coordinately at sites of envelope-spindle intersection. Both dismantling the spindle and NE fusion depend on membrane-remodeling ESCRT-III filaments and enzymes such as the microtubule-severing enzyme, SPASTIN, and the ESCRT-III disassemblase, VPS4^6,13,14^. The crucial nature of deploying this pathway with spatiotemporal precision is underscored by the discovery that deregulation of this pathway leads to genomic instability and nuclear malformation^6,13–16^. There are twelve human ESCRT-III proteins that polymerize on membrane surfaces to form membrane-remodeling filaments. Among these, CHMP7 is unique because it contains N-terminal VPS25-like winged helix domains, followed by a C-terminal ESCRT-III polymerization domain^17^. As part of the ESCRT-II complex, VPS25 acts to nucleate ESCRT-III filament formation^18,19^. Interestingly, CHMP7 acts upstream of other ESCRT-IIIs that function at the NE^6,11,13^. With CHMP7’s non-canonical ESCRT-II/III hybrid architecture, it is not known whether CHMP7 acts to polymerize other ESCRT-III proteins by its VPS25-like domains, whether it can polymerize itself, or whether it integrates both features. While it is known that the LEM-domain protein LEM2 interacts directly with CHMP7 and is required for its recruitment to the nuclear envelope^11,12^, it is unclear how the ESCRT pathway is activated at specific sites where spindle disassembly must take place in coordination with nuclear membrane remodeling. Our investigation of these questions has revealed a new paradigm for nuclear envelope closure in which unexpected properties of LEM2, including the ability to undergo phase separation and to co-polymerize with CHMP7, are key.

## LEM2 forms liquid droplets for core localization

Using live cell imaging, we observed that full-length LEM2 transiently concentrated at the NE core and colocalized with BAF as nuclear transport cargo began accumulating (Fig. 1A, Extended Data Fig. 1A-C, Supplementary Table 1). The LEM domain of LEM2 bound BAF in vitro, and this interaction was necessary for LEM2 targeting (Fig. 1A, Extended Data Fig. 1C-E). Specifically, a four-amino acid (AA) substitution in the LEM domain (LEM2_m21_), analogous to the m24 mutation described for the related LEM family protein Emerin that precludes BAF binding^20^, disrupted LEM2 concentration at the core during this window.

**Fig. 1.**
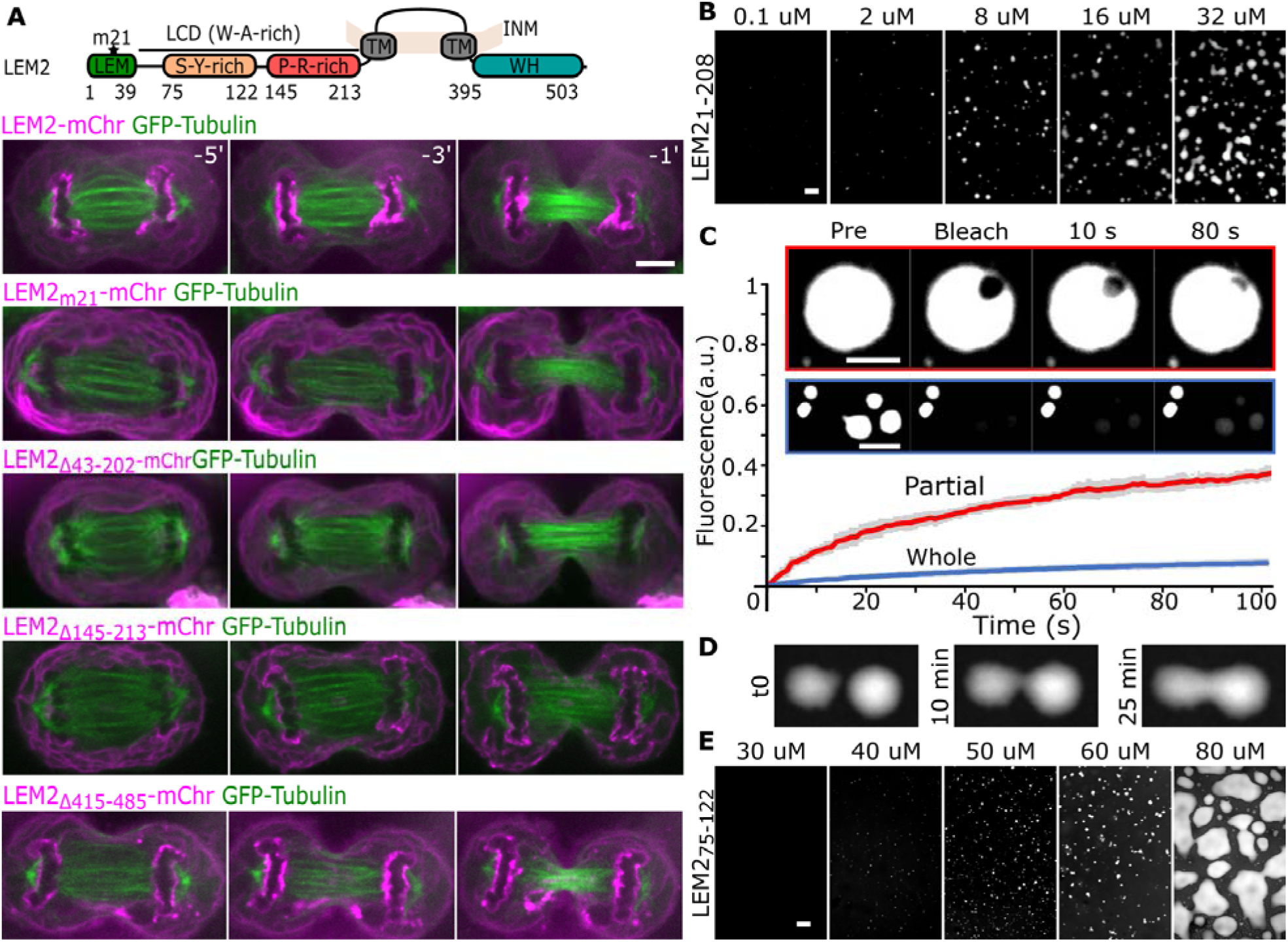
A low-complexity domain within LEM2 forms liquid droplets and is required for core localization. (A) Top: domain architecture of LEM2. Bottom: Live cell localization of LEM2-mChr and indicated deletions and mutations. Time 0 refers to full cleavage furrow ingression in late anaphase. In the m21 mutant, AA 21-24 are substituted with A. (B) Concentration-dependent droplet formation of LEM2_1-208_. (C) Whole and partial droplet fluorescence recovery after photobleaching (FRAP). N=3, bars are standard deviations (SD). (D) Real-time fluorescence imaging of LEM2_1-208_-droplet fusion. (E) Concentration-dependent droplet formation of S-Y rich peptide. Scale 5 µm (A-E).

Examination of LEM2’s sequence reveals an extended low complexity domain (LCD, AA 40-213). LCDs can mediate phase separation via multivalent, low-affinity interactions that scaffold segregated, liquid biochemical environments^21,22^. Deletion of LEM2’s LCD (LEM2_Δ43-202_) disrupted LEM2 concentration at the core despite the presence of the LEM domain, indicating that the LEM domain and the LCD are each necessary but neither is sufficient for localization (Fig. 1A). LEM2’s LCD diverges from the average protein’s amino acid composition with increased Alanine (A) and Tryptophan (W) content, and two subdomains distinguished by elevated Serine (S) and Tyrosine (Y) or Arginine (R) and Proline (P) levels, respectively (Fig. 1A, Extended Data Fig. 1F-G). Other proteins with comparable compositional biases form phase-separated liquid droplets^22,23^. To determine whether LEM2’s LCD undergoes phase separation, we first analyzed the behavior of purified extra-lumenal domains of LEM2 in vitro. At physiological salt and pH, we observed that the entire N-terminal domain, which includes the LEM motif and LCD (LEM2_1-208_), spontaneously formed spherical droplets that displayed liquid-like properties: droplet fusion and inter- and intra-droplet diffusion (Fig.1B-D, Extended Data Movies 1 and 2). Interestingly, at higher concentrations, LEM2_1-208_ droplets also spread out to wet the glass surface, indicative of low surface tension (Fig.1B). Analyzing short, synthetic peptides tiled across LEM2’s LCD sequence revealed that the S-Y-rich region^23^ of LEM2’s LCD was sufficient to form similar liquid droplets that retained this wetting behavior on glass (Fig. 1E, Extended Data Fig. 2A). In summary, we found that the LEM and LCD domains of LEM2 are each required for LEM2 enrichment at the spindle-containing core region of nascent nuclei through direct BAF binding and liquid-liquid phase separation, respectively.

## A liquid coat for spindle microtubules

To better understand how LEM2’s LCD mediates core enrichment, we performed stimulated emission depletion (STED) microscopy of immunostained HeLa cells. This imaging revealed that within the core region, endogenous LEM2 concentrates around spindle microtubules (MTs) near their interface with chromatin (Fig. 2A). In vitro analysis showed that the LEM2_1-208_ droplets have the capacity to up-concentrate tubulin, and nucleated stabilized MT from soluble tubulin, at physiological salt and pH conditions that were otherwise prohibitive for tubulin polymerization in the absence of LEM2_1-208_ (Extended Data Fig. 2B). Furthermore, LEM2_1-208_ droplets bound to MTs and bundles of MTs, wetting the MT surface and coating them completely (Fig. 2B). As expected for a liquid coating, LEM2_1-208_ found on MTs and MT bundles recovered its fluorescence after photobleaching, by contrast with the underlying MT lattice (Fig. 2B-C).

**Fig. 2.**
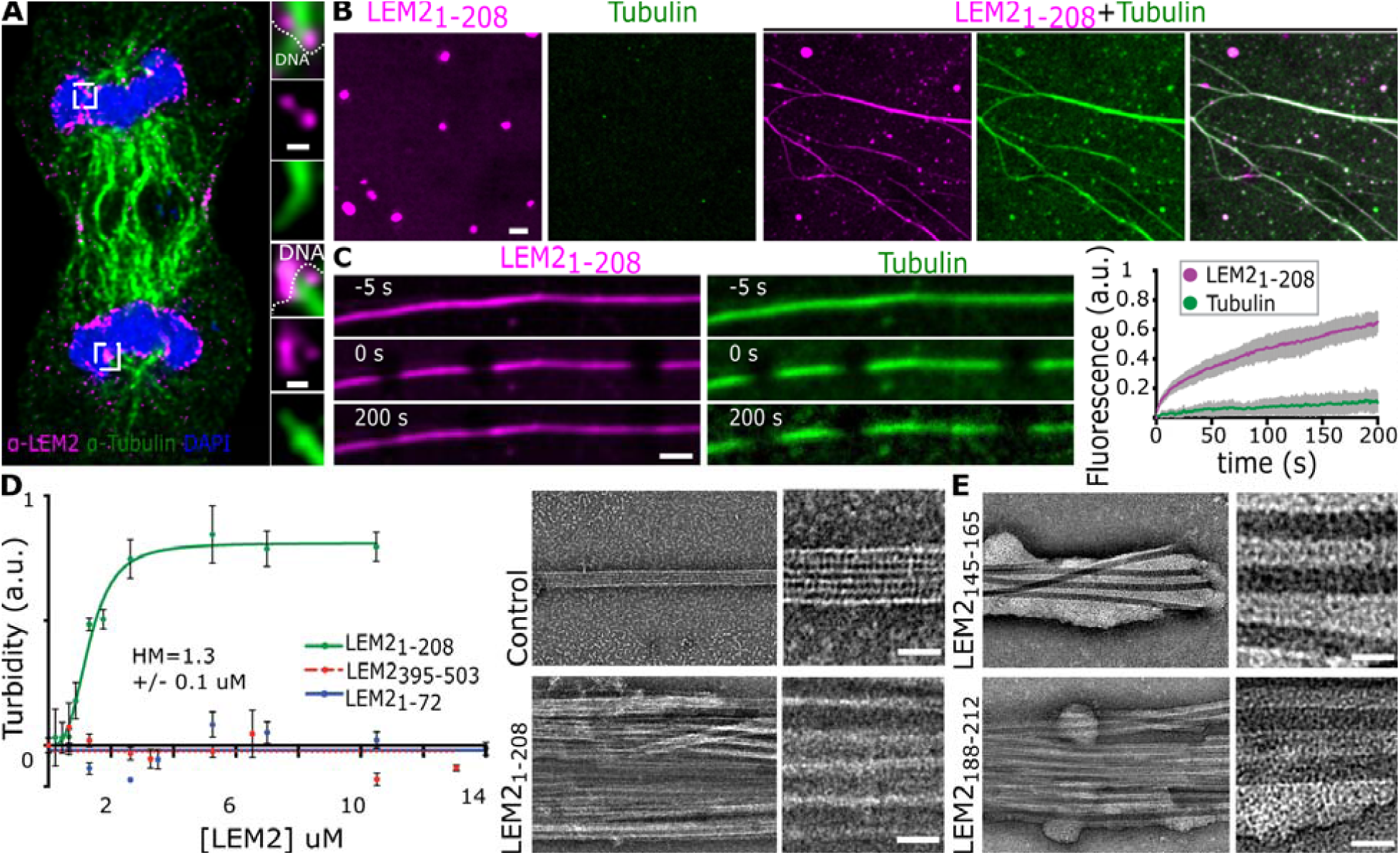
LEM2’s low complexity domain forms a liquid protein “coating” around spindle microtubules, stabilizing bundles. (A) STED imaging of endogenous LEM2 in late anaphase. Scale 150 nm. (B) LEM2_1-208_ droplets alone or mixed with MTs. Scale 2 µm. (C) FRAP of LEM2_1-208_-coated MT bundles. N=17, bars are SD, Scale 1 µm. (D) Light scattering-based quantification of MT bundling b indicated constructs, bars are standard error of the mean (left) and electron micrographs showing LEM2_1–208-_ -coated MT bundles (right). Half Maximum (HM) (E) Electron micrographs showing MT bundles induced by P-R-rich sequences from the LCD domain of LEM2. Scale 25 nm (D-E).

A light-scattering assay for MT-bundle formation, corroborated by negative stain electron microscopy (EM), revealed that the LEM2 LCD was required for the observed MT-bundling activity. Truncated protein constructs missing the LCD (LEM2_395-503_, LEM2_1-72_) did not promote the formation of light-scattering MT bundles in vitro. Furthermore, MT bundle formation by LEM2_1-208_ was concentration-dependent and saturable, with a half-maximal scattering concentration of 1.3 µM LEM2_1-208_ (Fig. 2D, Extended Data Fig. 2C). Negatively stained MT bundles detected by EM lost their characteristic tubulin surface fine structure, occluded by the LEM2_1-208_ coating (Fig. 2D-E). Using synthetic peptides that span LEM2’s LCD, we identified two separate sub-motifs within the P-R-rich region that were sufficient to induce MT bundling (Fig. 2E, Extended Data Fig. 2A, D-G). Importantly, this P-R-rich region was also required for LEM2 core localization in living cells (Fig. 1A). In summary, the LCD’s propensity for phase separation and its affinity for microtubules govern LEM2’s localization to spindle microtubules during NE reformation, with the LEM domain further restricting this to the surface of chromatin disks.

## LEM2 and CHMP7 initiate sealing at the reforming NE

We previously reported that the C-terminal winged helix (WH) domain of LEM2, also referred to as the MSC domain^11,24^, directly binds to CHMP7 in vitro ^11^. Testing the functional consequence of this binding interaction in vivo, we found that expression of a construct missing a portion of the WH-domain (LEM2_Δ415-485_-mChr) localized correctly to the core but was defective in GFP-CHMP7 recruitment when compared to the expression of full-length LEM2-mChr (Fig. 1A and 3A). This phenotype was dominant negative, as it was observed despite the presence of endogenous LEM2. Furthermore, overexpression of siRNA resistant LEM2_Δ415-485_-mChr following depletion of endogenous LEM2 failed to rescue recruitment of IST1, an ESCRT-III protein known to be downstream of LEM2 and CHMP7 (Fig. 3B and C)^6,11^. In contrast, overexpression of siRNA resistant full-length LEM2 notably enhanced recruitment of endogenous IST1 to the nascent nuclear envelope, reflected by both elevated IST1 levels and premature recruitment (Fig. 3C, Extended Data Fig. 3A-C, Supplementary Table 1). Together, these observations indicate that the C-terminal WH-domain of LEM2 is necessary and sufficient to recruit CHMP7 and downstream ESCRT proteins to reforming nuclear membranes.

**Fig. 3.**
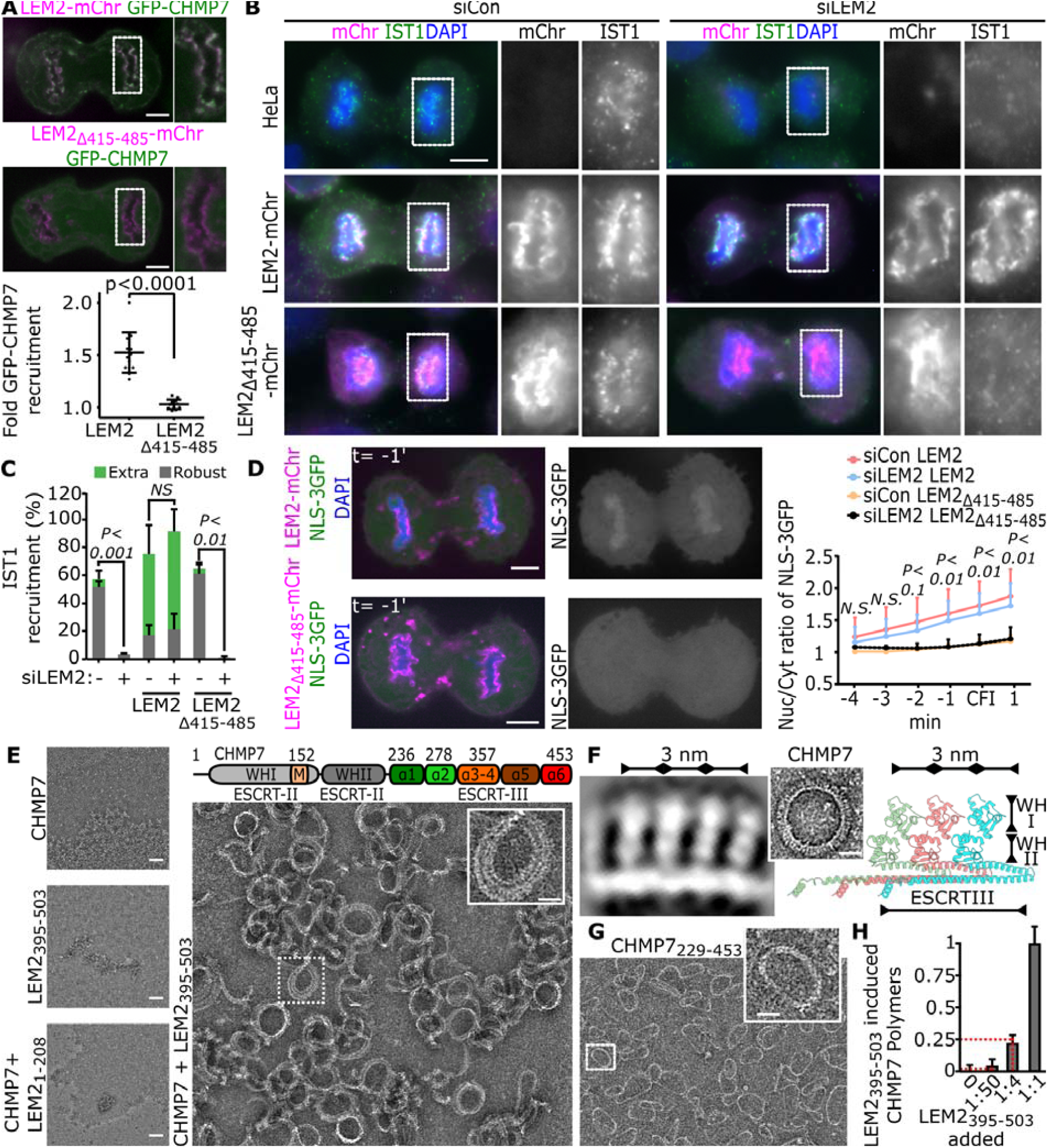
LEM2’s C-terminal WH domain induces CHMP7 polymerization and is required to seal the nascent NE. (A) Fold enrichment of GFP-CHMP7 at sites where full-length LEM2-mChr or LEM2_Δ415-485_-mChr concentrates at the NE in late anaphase. Bars are SD. (B) Recruitment of endogenous IST1 with indicated siRNA treatments and expression of siRNA resistant constructs. (C) Quantification of robust and extra-robust IST1 recruitment to chromatin disks in late anaphase. Error bars are standard error of the mean. Not significant (NS). (D) Nuclear import of NLS-3xGFP with indicated siRNA treatments and expression of siRNA resistant constructs. Bars are SD. (E) CHMP7 polymerization assay with indicated components assayed by negative stain EM. (F) Left: High magnification 2D class average of polymerized CHMP7, Low-mag polymerized ring of full-length CHMP7 (inset). Right: Homology model of polymeric CHMP7. (G) Negative stain EM of indicated construct. (H) SDS-page based relative quantification of polymerized and pelleted CHMP7 with different ratios of LEM2_395-503_ present. Red lines indicate expected fraction of CHMP7 in the pelleted polymer, assuming 1:1 stoichiometric polymer. N=3. Error bars are SD. Scale 10 µm (A-D) 50 nm (E) 20 nm for insets (E-H).

To test the biological role of LEM2’s WH-domain in establishing nuclear compartmentalization, we measured initial stages of NLS cargo nuclear accumulation. We found that NLS-3xGFP began to accumulate prior to cleavage furrow ingression when full-length LEM2-mChr was overexpressed (Fig. 3D, Extended Data Fig. 3D). In contrast, the initiation of NLS-cargo nuclear accumulation was abrogated by expression of LEM2_Δ415-485_-mChr, indicating that LEM2 works directly with CHMP7 to promote nuclear compartmentalization during early anaphase (Fig. 3D, Extended Data Fig. 3D).

## LEM2 activates CHMP7 to form looping copolymers

To elucidate how LEM2 engages CHMP7 for early nuclear compartmentalization and ESCRT-III recruitment, we turned to in vitro approaches. Full-length human CHMP7 purified as a stable monomer, and incubation of CHMP7 with LEM2_1-208_ had no discernable effect on CHMP7’s oligomeric state. Incubation of CHMP7 with the C-terminal LEM2_395-503_ WH domain, however, triggered the assembly of multi-layered, looping polymers with an average inner diameter of ∼50 nm (Fig. 3E). Liposomes were also sufficient to trigger polymerization of full-length CHMP7, with many polymers bound to the membrane surface (Extended Data Fig. 3E-F). 2D alignment and image averaging of these membrane-induced polymers, which were in a favorable orientation for subsequent classification, revealed a repeating polymeric unit comprised of a continuous polymeric strand studded with repeating perpendicular spikes (Fig. 3F, Extended Data Fig. 3G). The dimensions of the periodic features in this polymer match those of the homologous structure of “open” conformation human CHMP1B and yeast Snf7 polymers, suggestive of an ESCRT-III polymeric strand and protruding ESCRT-II-like tandem WH-domains^19,25,26^ (Fig. 3F, Extended Data Fig. 3G, Extended Data Table 1). A truncated CHMP7 fragment (CHMP7_229-453_) comprised only of the ESCRT-III domain spontaneously polymerized into rings during purification, also with an average diameter of ∼50nm, but devoid of perpendicular spikes (Fig. 3G). The CHMP7 ESCRT-III domain-only polymers confirm the domain arrangement model of the full-length CHMP7 polymer, and are consistent with an autoinhibitory function for CHMP7’s N-terminal WH domains. To probe the mechanism of LEM2-mediated activation of CHMP7 polymerization, we quantified CHMP7 polymerization with increasing concentrations of LEM2_395-503_ and found that LEM2’s WH domain induced a stoichiometric concentration of CHMP7 to polymerize, consistent with activation by co-polymerization rather than by nucleation (Fig. 3H). Together, these data show that CHMP7 can polymerize into looping strands with a polymerizing ESCRT-III domain regulated, in part, by autoinhibitory VPS25-like domains. Interestingly, CHMP7 polymerization is both activated and stoichiometrically limited by LEM2_395-503_.

### A domain replacement mechanism controls CHMP7

To investigate the mechanism of CHMP7 autoinhibition and release by LEM2’s WH-domain, we employed a quantitative isotopic labeling and cross-linking mass spectrometry (XL-MS) approach (Fig. 4A, Supplementary Table 2). Labeling monomeric CHMP7 with a heavy crosslinker and LEM2-induced polymers with a light crosslinker, we quantitatively compared crosslink enrichment between samples. We identified 24 cross-links specific to monomeric CHMP7. Hybrid peptide mapping revealed that the N- and C-termini of CHMP7 fold back to interact with each other and with the α1-α3 helices of CHMP7’s ESCRT-III domain (Fig. 4A-B, and Extended Data 4A-E). These interactions were significantly reduced or absent in polymerized CHMP7 incubated with the activating and co-assembling LEM2 WH-domain (Fig. 4A-B, Extended Data Fig. 4 and Extended Data Table 1), consistent with a conformational change from a closed, monomeric state into an open, polymeric state, as corroborated by EM visualization (Fig. 3E). In total, 19 cross-links were enriched in the polymeric sample, with interactions between α1-α3 of CHMP7 and LEM2_395-503_. Polymer-specific cross-links between N-terminal amines of neighboring CHMP7 molecules, and between adjacent LEM2_395-503_ molecules, are consistent with the co-assembly of both proteins (Fig. 4A-B, Extended Data Fig. 4). Mutating conserved residues implicated by XL-MS, within CHMP7 α1-3, impaired LEM2_395-503_-induced CHMP7 polymerization (Fig. 4C and D, Extended Data Fig. 4F-G). These data suggest that autoinhibited CHMP7 is activated by LEM2_395-503_ binding to the α1-3 region of CHMP7, and this, in turn, enables copolymerization. In the copolymer conformation, CHMP7’s N-terminal WH-domains are presumably displaced from interactions with the ESCRT-III domain and may thus become available for downstream ESCRT-III binding (Fig. 3F, Fig. 4D).

**Fig. 4.**
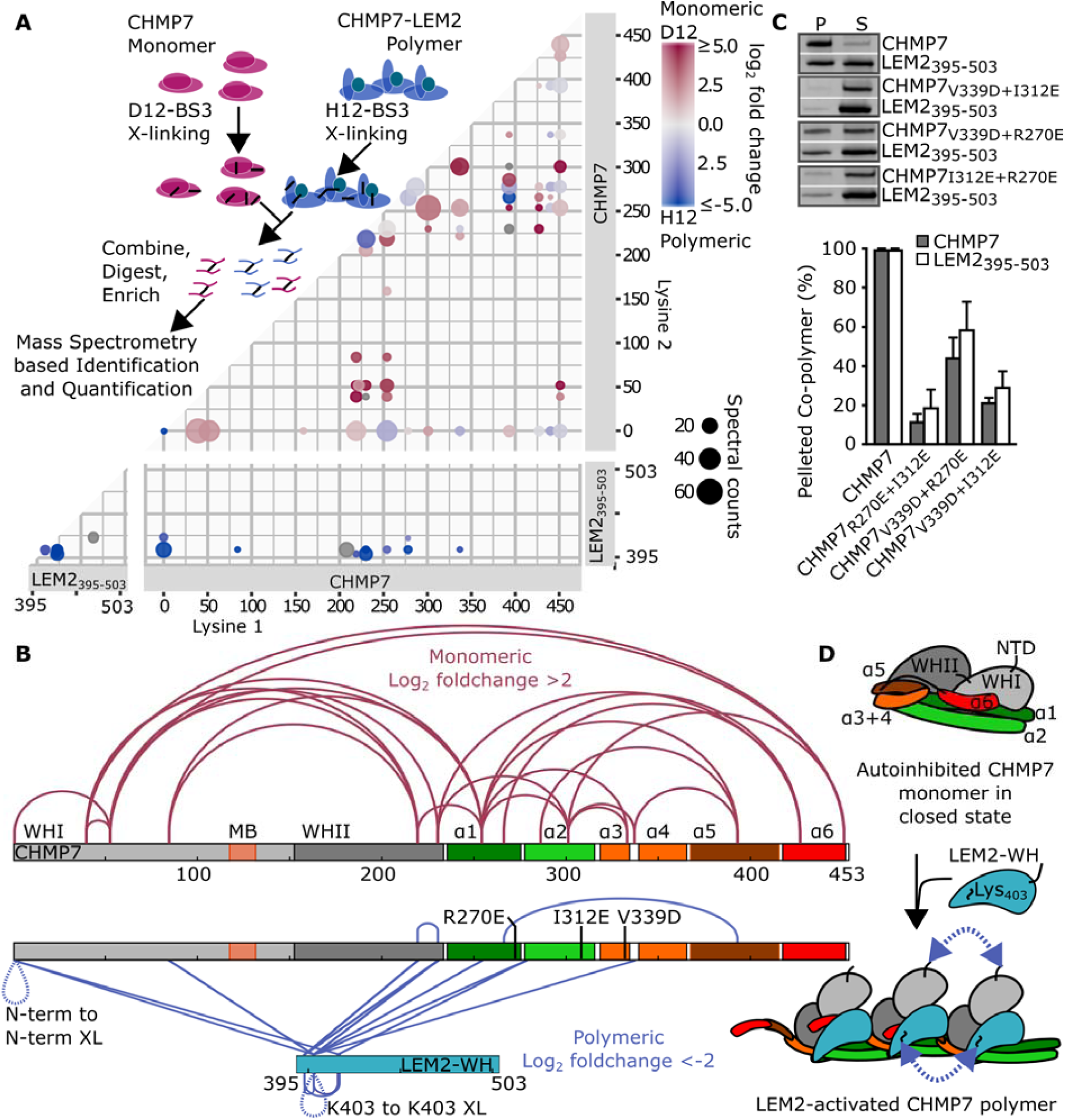
The WH domain of LEM2 activates CHMP7 polymerization by binding the ESCRT-III-core domain to relieve CHMP7 autoinhibition. (A) Workflow of lysine-lysine hybrid peptide mapping using XL-MS (top) and results represented by a domain interaction matrix (bottom). BS3 cross-links surface accessible Lys residues with C_α_-C_α_ distances <∼3nm. Each dot represents a pair of crosslinked Lys-Ly residues with area scaled to the sum of spectral counts and the color indicating the enrichment of each interaction in either monomeric CHMP7 (red) or LEM2_395-503_-CHMP7 polymer (blue). (B) Interaction that were enriched more than 4-fold are mapped on to the protein primary structure. Tested mutations are indicated. MB, membrane binding. (C) Top: SDS-PAGE of protein in the pellet (P) or supernatant (S) following centrifugation of LEM2_395-503_ incubated with wild type or mutant CHMP7. Bottom: Recovery in the pellet. Bars are SD of n=3. (D) Cartoon summary of LEM2-mediated CHMP7 activation. Arrows illustrate the interaction of neighboring molecules identified by XL-MS.

### NE reformation depends on the LEM2 pathway

To understand the biological consequences of LEM2 copolymerization with CHMP7 in nuclear envelope remodeling and spindle disassembly, we studied nuclear phenotypes in cells following endogenous LEM2 depletion. Previous work has shown that siRNA depletion of LEM2 causes a nuclear deformation phenotype in interphase, and is lethal with extended time^27^. To address when nuclear defects arise, we synchronized cells following LEM2 depletion and imaged their progression from anaphase to late telophase. While LEM2-depleted cells progressed to anaphase without noticeable defects, a strong nuclear and tubulin-morphology phenotype began to emerge in late anaphase and persisted past telophase (Fig. 5A-C, Extended Data Fig. 5A-B, Movie 3). Depletion of LEM2 resulted in severely misshapen nuclei and 3D imaging revealed thick MT bundles piercing through the nucleus in a channel lined by nuclear membrane (Fig. 5B, Extended Data Fig. 5). The significance of this nuclear morphology phenotype is underscored by the appearance of DNA damage—a hallmark of defective nuclear integrity—in newly-formed nuclei (Fig. 5D, Extended Data Fig. 5C).

**Fig. 5.**
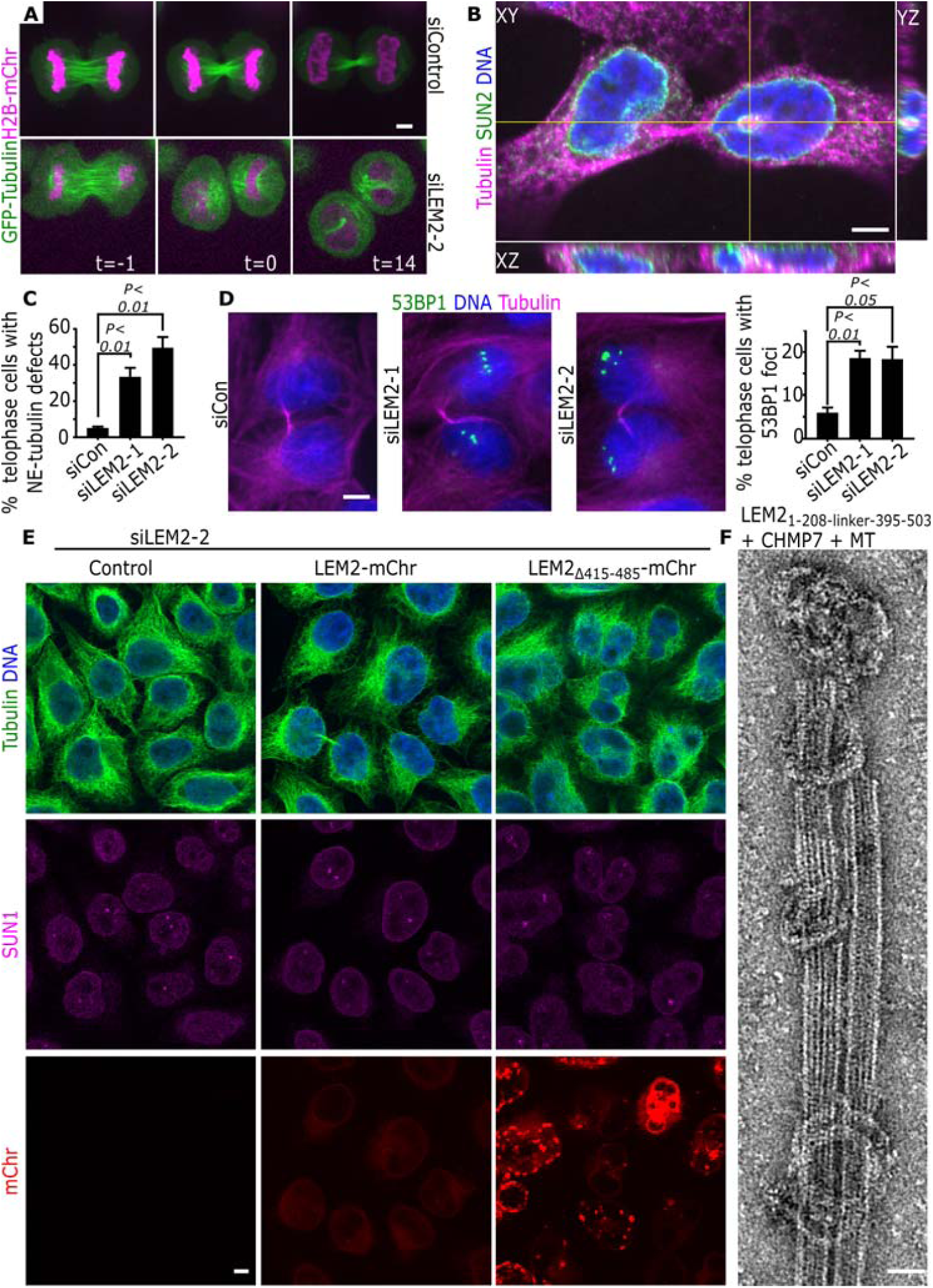
LEM2 and ESCRT-III are essential for spindle disassembly and nuclear integrity. (A) Live-cell imaging of GFP-Tubulin in siRNA treated cells. Time 0 refers to cleavage furrow ingression in late anaphase. (B) Orthogonal view of a representative telophase cell following treatment with siLEM2. (C) Tubulin defect scoring. Error bars are standard error of the mean. (D) 53BP1 localization in telophase U2OS cells following siRNA treatment. Graph show percent of telophase cells with ≥5 53BP1 foci. (E) HeLa cells after siRNA treatment and rescue construct expression. (F) Negative stain EM of co-incubated indicated proteins. Scale 5 µm (A-E) 25 nm (F).

Interestingly, depletion of endogenous LEM2 alongside expression of LEM2_Δ415-485_-mChr proved more deleterious to nuclear morphology at interphase than simply depleting LEM2 (Fig. 5E, Extended Data Fig. 5D). Since the N-terminal region of LEM2_Δ415-485_-mChr is intact, while the WH domain is not, this observation highlights that focal enrichment at the spindle-NE-chromatin interface, a function of LEM2’s N-terminus, must be balanced by downstream ESCRT-III activity, a function of LEM2’s C-terminus. These activities were similarly integrated in vitro, where a construct bearing the extra-lumenal domains of LEM2 connected by a 14-AA linker, LEM2_1-208-LINKER-395-503_, induced the formation of CHMP7 polymers looping at and around small MT bundles (Fig. 5F).

In this work we explain the molecular mechanisms underlying LEM2’s function as a site-specific adaptor for mitotic ESCRT-III functions at the NE. Two distinct features specify the localization of LEM2 to sites along the nascent NE that are destined for coordinated membrane fusion and spindle disassembly: LEM domain interactions with BAF and its low complexity domain, which is responsible for liquid-liquid phase separation and wetting the MT-chromatin interface. As an integral inner nuclear membrane protein, LEM2 tethers the reforming NE to the chromatin surface and to the spindle, simultaneously recruiting the ESCRT-II/ESCRT-III chimeric protein CHMP7 with its C-terminal WH-domain. We further reveal how LEM2 initiates assembly of the ESCRT-III complex through direct activation of CHMP7. Together LEM2 and CHMP7 copolymerize and loop around spindle MTs, creating a molecular “O-ring” to promote nuclear compartmentalization, even before the spindle is fully disassembled (Extended Data Fig. 1A-B, Fig. 3D, Fig. 6).

**Fig. 6.**
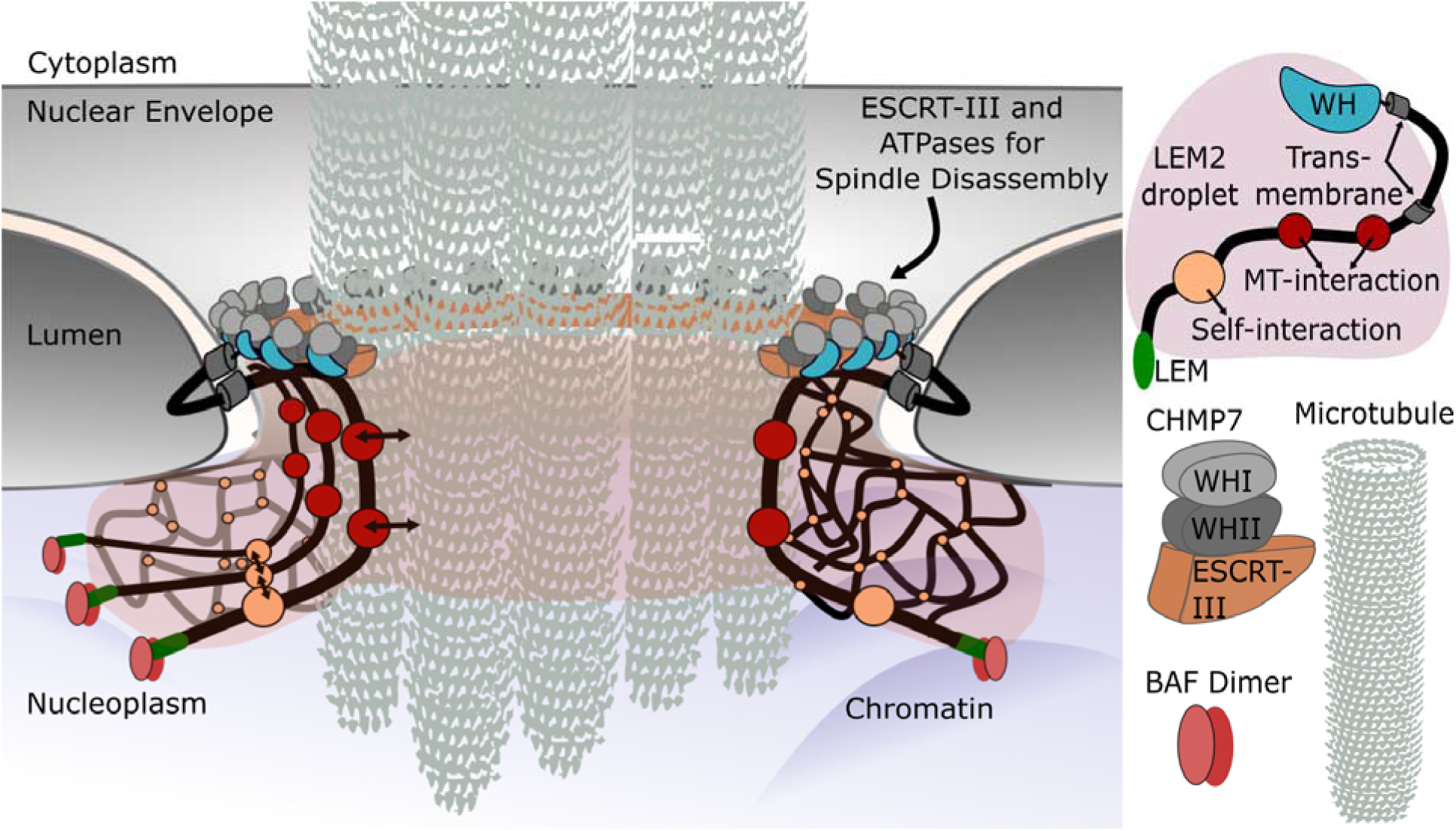
Model of a molecular O-ring formed by LEM2 and CHMP7 during nuclear reformation that is required for spindle disassembly and to seal the nascent NE. A deformable LEM2 phase forms around the spindle which is anchored to chromatin surface by LEM2s interaction with BAF. LEM2 regulates ESCRT recruitment by inducing CHMP7 polymerization for nuclear membrane remodeling and sealing.

Our data suggest that the polymeric surface of spindle MTs nucleates LEM2 condensation. We propose that the deformable wetting behavior of LEM2 on the surface of MT bundles may serve as a fluid coating at the junction where coalescing membranes encounter bundles of residual microtubules. As LEM2 is embedded in the NE of cells, wetting of MT bundles in vivo may perform mechanical work by restricting the size of gaps in the NE and drawing the membrane into close apposition with MT bundles before ESCRT-III constriction and fusion of the membrane. We propose that during nuclear envelope reformation LEM2 constitutes a deformable liquid phase between NE and spindle MT, surrounding spindle MT bundles in a circular cross-section. This flexible toroid of LEM2 recruits and polymerizes with CHMP7, in a LEM2-CHMP7 interaction that is required to initiate nuclear compartmentalization. CHMP7, too, has a physical connection with the membrane at this junction ^28^. Thus, we propose that the LEM2-CHMP7 assembly provides a pliable circular connection between membrane and MT, pre-sealing the NE prior to complete spindle disassembly and membrane fusion by the ESCRT-III pathway. These results further raise the intriguing possibility that liquid-liquid phase separation of transmembrane or membrane-associated proteins provides important properties that may contribute to membrane remodeling activities more generally^29^.

## Acknowledgments

We thank Wesley I. Sundquist and Sy Redding for critically reading the manuscript. For reagents, technical advice, and discussions we thank the Frost and Ullman labs, as well as the Nikon Imaging Center at UCSF. We thank Lauren Williams for helping score and the lab of Michelle Mendoza for generously sharing microscope resources. We also thank the UCSF Center for Advanced cryoEM, including Alexander Myasnikov, David Bulkley and Michael Braunfeld.

## Author contributions

Author contributions: A.v.A, D.L., I.E.J., M.T., A.L.B., K.S.U., and A.F. designed research; A.v.A, D.L., I.E.J., M.T., and S.M.P. performed research; A.v.A, D.L., I.E.J., M.T., K.S.U., and A.F. analyzed data; and A.v.A, D.L., I.E.J., K.S.U., and A.F. wrote the paper.

## Extended Data information titles and legends

**Extended Data Figure 1.**
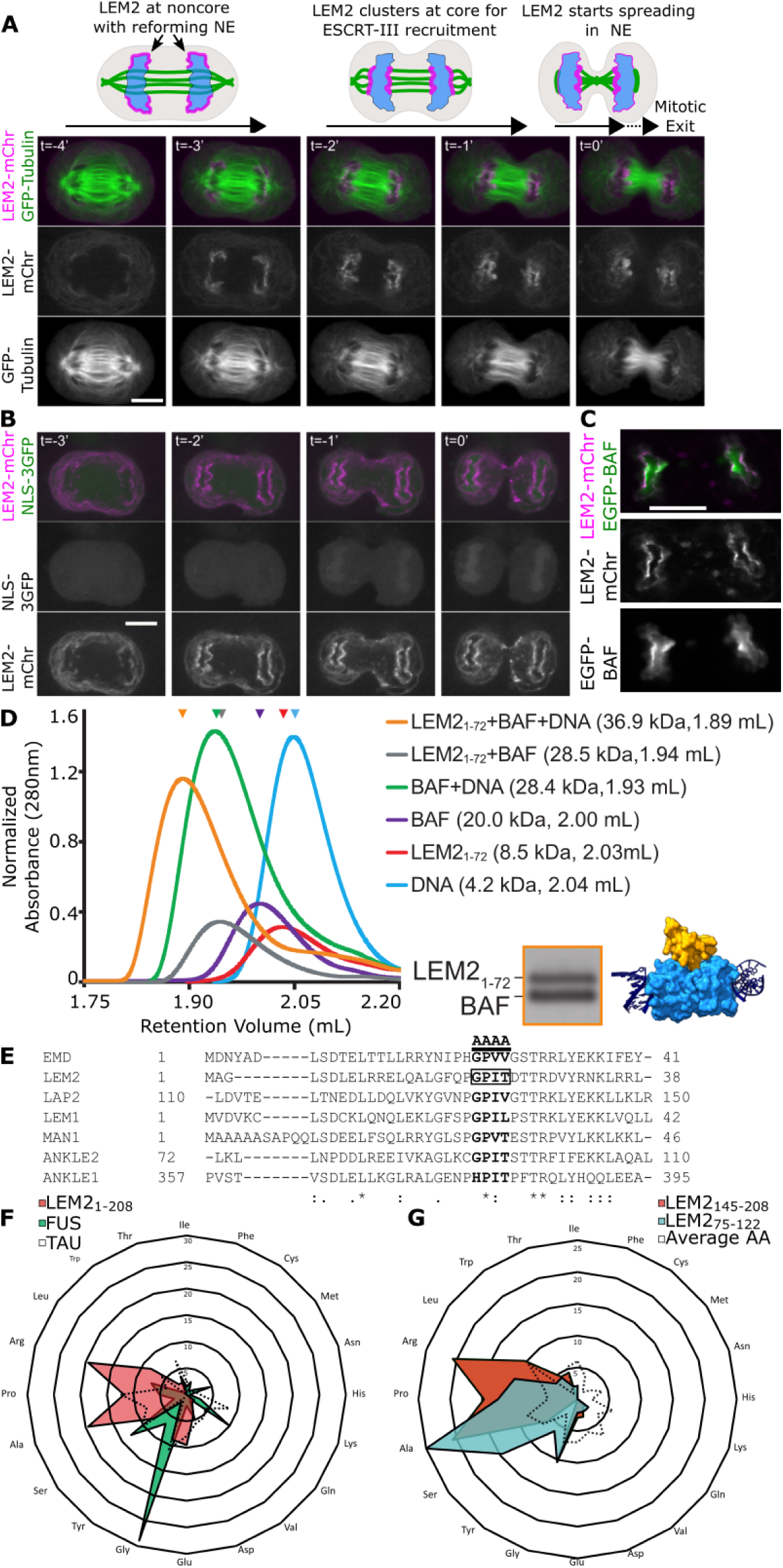
LEM2 dynamically colocalizes at the core with BAF. (A) Live imaging of HeLa cells stably expressing LEM2-mCherry and GFP-Tubulin. The cartoon summarizes LEM2 localization through anaphase. Scale bar 10 µm. (B) HeLa cells stably expressing NLS-3xGFP and transiently expressing LEM2-mCherry were live-imaged. Zero is the time of complete cleavage furrow ingression (min). Scale bar 10 µm. (C) HeLa cells stably expressing LEM2-mCherry and EGFP-BAF were live-imaged in anaphase. Scale bar 10 µm.(D) Absorbance 280 nm as a function of retention volume (mL) from analytical size exclusion chromatography. Retention volumes for major peaks (arrowheads) and predicted molecular weights for protein or protein-DNA complexes are listed. Lower right: A homology model for LEM2_1-72_ -BAF-DNA complex, based on high-resolution interfaces of BAF with DNA (2BZF), and BAF with LEM-domain of EMERIN (2ODG) (*36*, 5*3*). (E) Sequence alignment of LEM domains across LEM family proteins showing conservation of a four-amino acid (AA) sequence that, when mutated in EMD, disrupts BAF binding. Based on this alignment, the m21 mutation in LEM2 is analogous to the m24 mutation in EMD (*15*). (F) LEM2 N-terminus (LEM2_1-208_) percent amino acid composition compared to full length FUS and TAU proteins. (G) The LEM2 LCD shows elevated A and W levels compared to the average protein and has distinct S-Y-rich (LEM2_75-122_) and a R-P-rich (LEM2_145-208_) subdomains.

**Extended Data Figure 2.**
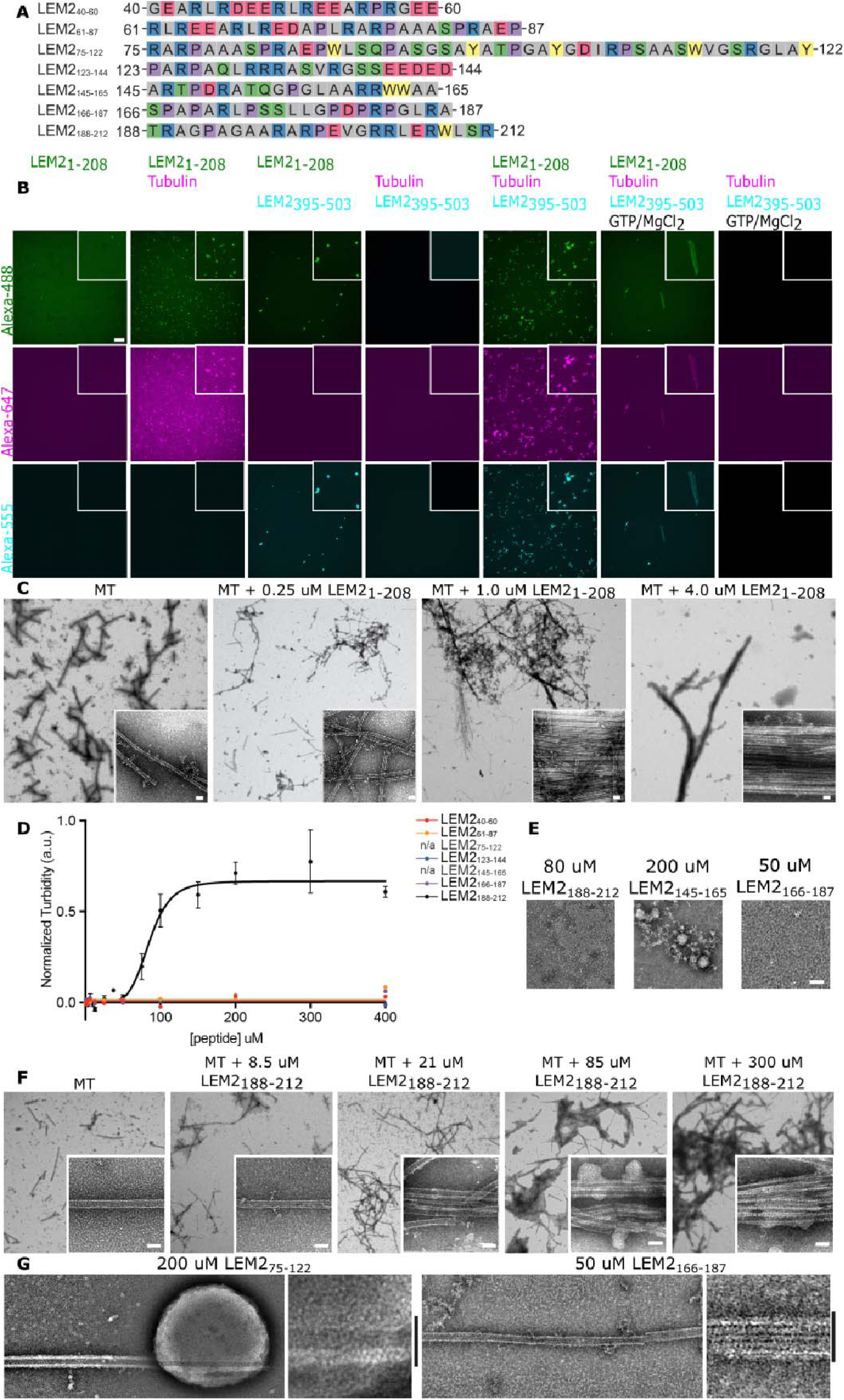
Microtubule bundling by LEM2-LCD reveals distinct droplet-forming and MT-bundling domains. (A) Amino acid sequences of 7 consecutive LEM2 peptides tiling the LCD. (B) Indicated proteins, 8 µM each, were incubated at physiological salt and pH. LEM2 was supplemented with 0.5 µM Alexa488-labeled His_6_-SUMO-LEM2_1-208_ for visualization. Scale bar is 10 µm. (C) Negative stain EM of MTs or MTs with different concentrations of LEM2_1-208_, corresponding to light scattering reactions. Scale bar 25 nm. (D) Light scattering-based quantification of MT bundling by indicated LEM2 peptides. Half maximal concentration of LEM2_188-212_ is 85.11+/-3.490 µM. LEM2_75-122_ and LEM2_145-165_ peptides were turbid in the absence of MTs, and each induced the formation of clusters too big to be analyzed by this assay upon exposure to MTs. (E) Representative negative stain EM of soluble LEM2 peptides (LEM2_166-187_, LEM2_188-212_), and a light scattering LEM2 peptide (LEM2_145-165_). Scale bar is 50 nm. (F) Negative stain EM of MTs or MTs with different concentrations of LEM2_188-212_, corresponding to light scattering reactions. Scale bar 25 nm. (G) Negative stain EM of LEM2_75-122_ or LEM2_166-187_ with MTs. Scale bars 25 nm.

**Extended Data Figure 3.**
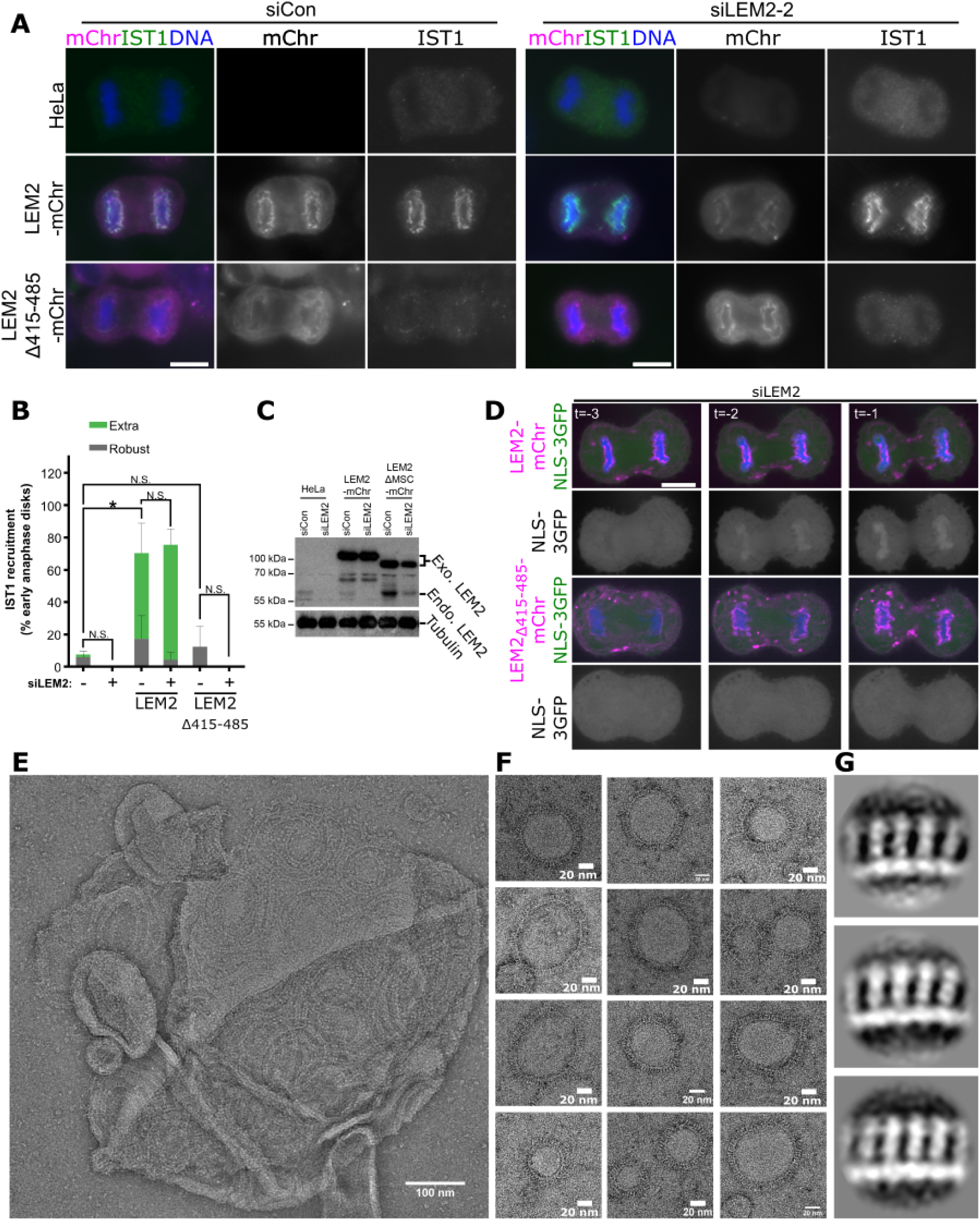
LEM2’s C-terminal WH domain is required for IST1 recruitment and nuclear compartmentalization. (A) Recruitment of endogenous IST1 with indicated siRNA treatment and expression of siRNA resistant constructs in early anaphase. Scale bars 10 um. (B) Quantification of the percent of early anaphase disks with robust and extra robust IST1 recruitment, as assessed by blind scoring. Bars represent the average of three independent experiments and are graphed showing standard error of the mean. Student’s t-test was used to determine p-values, which are indicated as follows: \**P* < 0.05; N.S., not significant. (C) Immunoblot showing overexpression of the siRNA-resistant full length LEM2-mCherry (84kDa) and LEM2_Δ415-485_-mChr (76kDa). Endogenous LEM2 is approximately 60kDa. (D) Representative examples of HeLa cells treated with siLEM2 and expressing the indicated siRNA-resistant LEM2 constructs and a nuclear localization signal fused to three GFPs (NLS-3xGFP), live-imaged through anaphase. Scale 10 um. (E) Negative stain EM of CHMP7 polymers covering a liposome. Scale bar 100 nm. (F) Negative stain EM of membrane-induced CHMP7 polymers used for class-averaging. Scale bar 20 nm. (G) Representative 2D-class averages from manually particles from polymers shown in F.

**Extended Data Figure 4.**
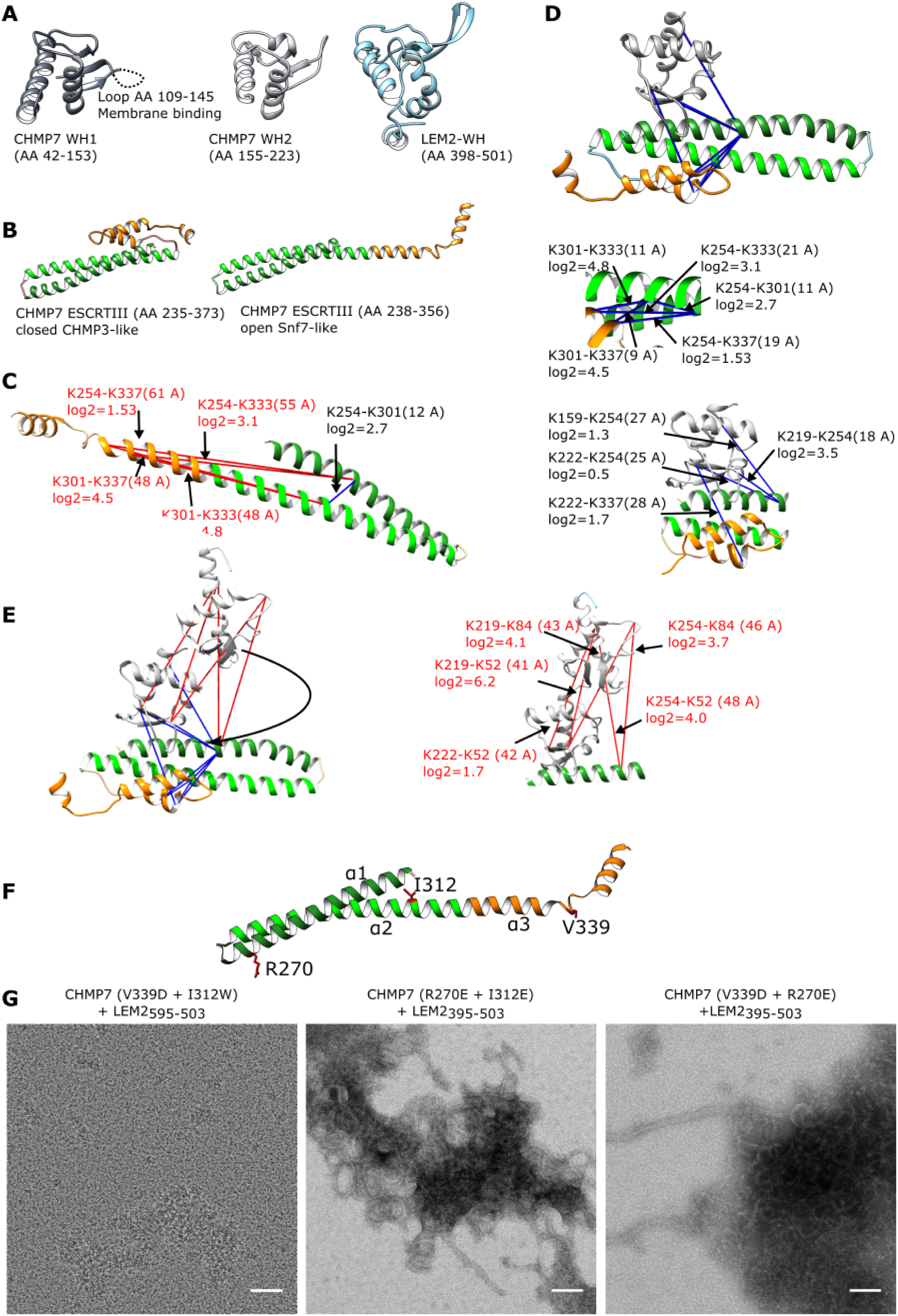
Homology modeling and crosslinking-mass spectrometry (XL-MS) reveal structural dynamics of CHMP7. (A) Homology models for the winged helix (WH) domains of CHMP7 and LEM2. WH1 of CHMP7 contains a membrane binding region indicated as a loop (*13*). (B) Homology models of the CHMP7 ESCRTIII-fold in open and closed confirmations. (C) Distance restraints identified from XL-MS analysis of the CHMP7 monomer were mapped to open and closed homology models, and the open ESCRTIII conformation was rejected. Satisfied restraints are show in blue, violated restraints are shown in red. Cross-linked lysine residues are indicated together with log_2_ fold change as compared to the polymeric CHMP7-LEM2_395-503_ sample. Mapped C_α_-C_α_ distances > 3 nm are considered violations. (D) A crystallographic interphase between VPS25 and an ESCRTIII helix (*22*) resembles the CHMP7 WH2 interaction to the CHMP7 ESCRTIII domain. All cross-links are satisfied when mapped to the closed CHMP7-ESCRTIII model (middle and lower panel) and agree with domain connectivity. A subset of cross-links was not satisfied when mapping WH1 instead to same interface (data not shown). (E) The cross-linking data from monomeric CHMP7 suggests a hinge region between CHMP7 WH1 and WH2 that allows its WH1 to move closer to the ESCRTIII core as compared to the conformation of polymerized CHMP7 in the class averages. (F) CHMP7 point mutations are shown in an open ESCRTIII-fold, representing polymerized CHMP7. (G) CHMP7 mutants were dialyzed with His_6_-SUMO-LEM2_395-503_ in a 1:2 molar ratio, and any resulting polymer was enriched by centrifugation and analyzed by negative stain EM. CHMP7_V339 + I312E_ did not form polymers in the presence of His_6_-SUMO-LEM2_395-503_ during dialysis. Scale bars are 50 nm.

**Extended Data Figure 5.**
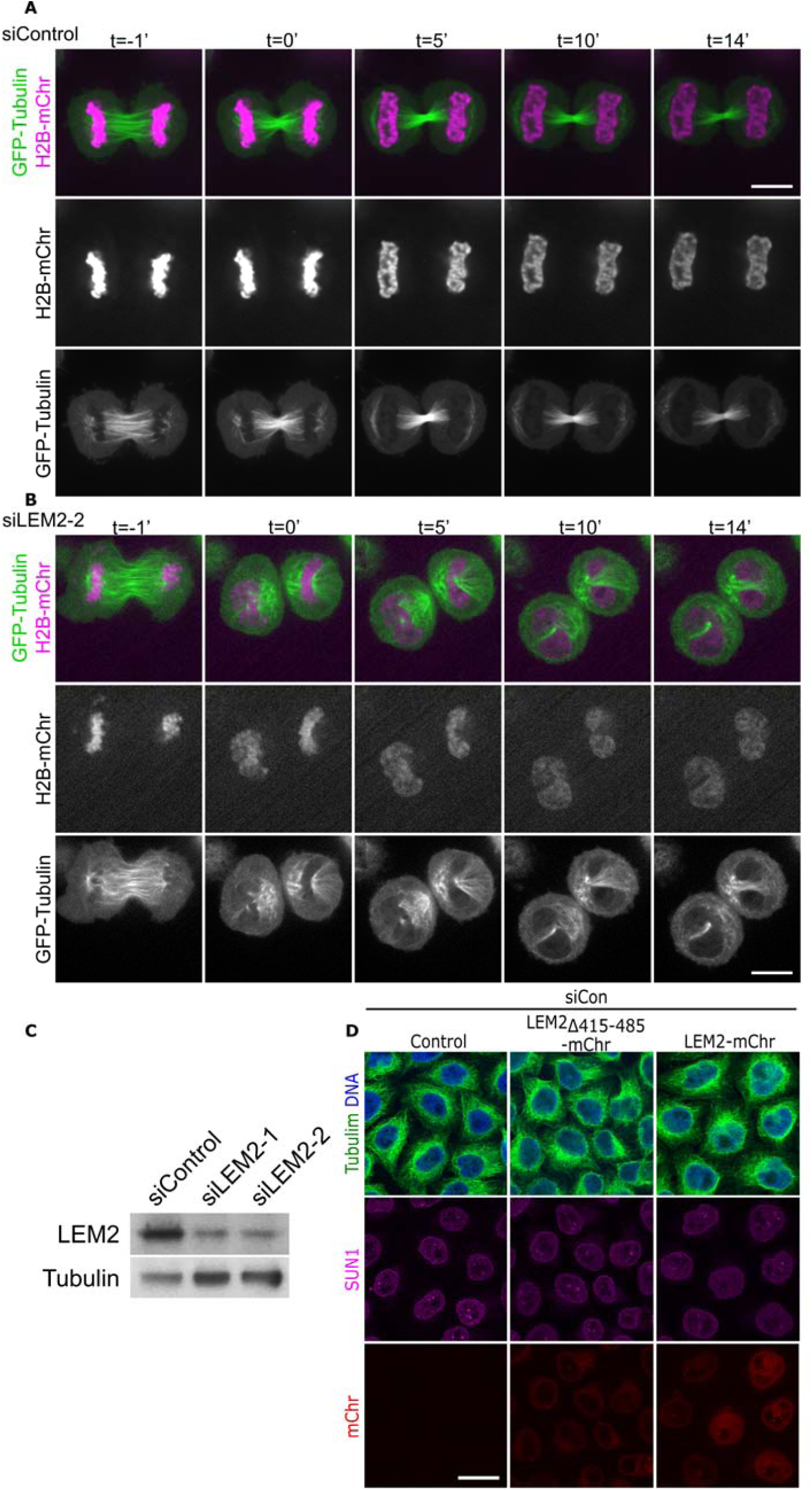
Development of post-mitotic tubulin phenotype with LEM2 depletion. (A-B) Live imaging through mitotic exit of HeLa cells stably expressing GFP-Tubulin and H2B-mChr with siRNA control (A) or siLEM2 treatment (B). (C) Immunoblot demonstrating LEM2 depletion by two independent siRNAs in U2OS cells. Scale bars 10 um. (D) Immunostained HeLa cells stably expressing no or indicated siRNA-resistant constructs were treated with control siRNA and imaged in interphase by spinning disk confocal microscopy. Scale bar 20 um.

## Methods

### Cytoskeletal proteins

Porcine brain Tubulin reagents were purchased from Cytoskeleton, Inc., including unpolymerized Tubulin (T238P), unpolymerized, HiLyte-647-labeled Tubulin (TL670M), and pre-formed microtubules (MT002).

#### Purification of His_6_-SUMO-tagged proteins

All purified proteins in this study were expressed in a pCA528 vector (WISP08-174; DNASU Plasmid Repository) in BL21-(DE3)-RIPL *Escherichia coli* cells using an N-terminal His_6_-SUMO affinity tag as described previously^11^. All plasmids listed in Table S2. Expression cultures (3- to 4-liter) were grown in ZY-5052 rich auto-induction media containing kanamycin, shaking (220 rpm) for 3 hours at 37°C, then overnight at 19°C. Cells were harvested by centrifugation.

#### Purification of BAF

Purification of full length, human BAF protein (Uniprot ID O75531) was adapted from a published protocol^5^. The purification was performed exactly as described, except His_6_-SUMO-BAF was cleaved with His_6_-Ulp1 protease (30 min, room temperature) to suit the use of the His_6_-SUMO affinity tag. Ultimately, Superdex 75 16/60 fractions containing BAF dimer were pooled, concentrated, and flash frozen in liquid nitrogen and stored as single use aliquots at −80°C.

#### Purification of LEM2 proteins

Protein constructs originating from the human LEMD2 protein (Uniprot ID Q8NC56), including LEM2_1-72,_ LEM2_1–208_, LEM2_395–503_, and LEM2_1-208-LINKER-395-503_ (linker sequence SAAGTGAGSGSAAS), LEM2_1-72_, LEM2_395–503_, and LEM2_1-208-LINKER-395-503_ constructs were purified as follows. Harvested cells were resuspended in lysis buffer (5 mL per gram cell pellet) containing 5% glycerol, 10 mM imidazole, pH 8.0, 2 μg/mL DNase1, lysozyme and protease inhibitors, with HEPES and KCl concentrations optimized individually for each LEM2 protein construct (20 mM HEPES, pH 7.0, 500 mM KCl for LEM2_1-72_ and LEM2_1-208-LINKER-395-503_, and 40 mM HEPES, pH 8.0, 350 mM KCl for LEM2_395–503_). Cells were lysed by sonication on ice, and clarified (10,000g, 30 min, 4°C). Clarified lysate was incubated with Ni-NTA agarose resin (Quiagen) (4 mL bed volume, 1.5 h, 4°C), washed extensively with lysis buffer, and protein was eluted with 5 column volumes of lysis buffer supplemented with 350 mM imidazole, pH 8.0. Eluate was spin-concentrated (Vivaspin 20, 3KDa MWCO, PES) to 5 mL, and dialyzed overnight at 4°C into storage buffer (20 – 40 mM HEPES, pH 7.0 – 8.0, 5% glycerol), with KCl concentrations optimized individually for construct (300 mM for LEM2_1-72,_ 500 mM for LEM2_1-208-LINKER 395-503, -_ and 350 mM for LEM2_395–503_). The His_6_-SUMO tag was cleaved by incubation with His_6_-Ulp1 protease during dialysis. LEM2 proteins were further purified by size exclusion chromatography using the Superdex 75 16/60 column (GE Life Sciences) in storage buffer, and LEM2-containing fractions were pooled, spin-concentrated (Vivaspin 6 3KDa MWCO, PES), and aliquoted.

LEM2_1-208_ was purified similarly, including phase separation as a preparative step. All steps were carried out at room temperature unless otherwise noted. Harvested cells were resuspended in lysis buffer (30 mM HEPES, pH 7.4, 500 mM KCl, 5% glycerol, 10 mM imidazole, pH 8.0, 2 μg/mL DNase1, lysozyme and protease inhibitors), and lysed by sonication with intermittent chilling on ice. Lysate was clarified by centrifugation (10,000g, 30 min), incubated with Ni-NTA agarose resin (4 mL bed volume, 1.5 h, room temperature), and washed extensively with lysis buffer. Protein was eluted with lysis buffer supplemented with 350 mM imidazole, pH 8.0. Droplet formation was induced to enrich for His_6_-SUMO-LEM2_1-208_: the imidazole eluate was diluted with ice-cold buffer (40 mM HEPES, pH 7.4, 5% glycerol) to drop the concentration of KCl to 50 mM. After 20 min incubation on ice, protein droplets were pelleted by centrifugation (10,000g, 10 min, 0°C), washed with 2 column volumes of ice-cold low-salt buffer (40 mM HEPES, pH 7.4, 50 mM KCl, 5% glycerol), and pelleted again. The pellet was resuspended in 5 mL of high-salt buffer (40 mM HEPES, pH 7.4, 500 mM KCl, 5% glycerol), to dissolve protein droplets for the remaining purification steps, and incubated with His_6_-Ulp1 protease (2 h). The mixture was incubated with Ni-NTA resin (2 mL, 1 h). Spin-concentrated, flow-through protein was further purified by size exclusion chromatography using the Superdex 75 16/60 column in high-salt buffer at 4 °C. Pooled, spin-concentrated, LEM2-containing fractions were dialyzed overnight into storage buffer (40 mM HEPES, pH 7.4, 350 mM KCl, 5% glycerol), spin-concentrated, and aliquoted.

Concentrated protein aliquots were snap-frozen in liquid nitrogen and stored at −80°C for future experiments. Aliquots were thawed and used only once. Final protein concentrations were 260 µM for LEM2_1-72_, ∼110 µM for LEM2_1–208_, 30 µM for LEM2_1-208-LINKER-395-503_, 14 µM for LEM2_395-503_ and 190 µM for His_6_-SUMO-LEM2_395-503_. Note that concentrations LEM2_395–503_ without the tag were limited, and LEM2_395–503_ was more stable with the His_6_-SUMO tag.

#### Purification of CHMP7 proteins

Purification of full-length, human CHMP7 (Uniprot ID Q8WUX9) and point mutants CHMP7_R270E + I312E_, CHMP7_R270E + V339D_, and CHMP7_I312E + V339D_ were adapted from previously published purifications^11^. The truncated CHMP7_229-453_ was purified differently. Harvested cells were resuspended in 5 mL ice-cold lysis buffer (40 mM HEPES, pH 8.0, 800 mM KCl, 5% glycerol, 10 mM imidazole, 2 μg/mL DNase1, 5 mM BME, protease inhibitors, and lysozyme) per gram cell pellet. Cells were lysed by sonication, on ice, and clarified (10,000g, 30 min, room temperature). Clarified CHMP7_229-453_ lysate was incubated with Ni-NTA agarose resin and spontaneously formed a gel composed of protein polymerized into rings, assayed by negative stain EM. His_6_-SUMO-CHMP7_229-435_ rings were eluted with imidazole (350 mM), and the eluate was collected with low-speed spin (1,000g) as a gel phase on top of the resin. The gel was scooped off, washed three times with buffer (40 mM HEPES, pH 8.0, 800 mM KCl, 5% glycerol, 1 mM DTT), and polymers were collected by centrifugation (5,000g) each time. The His_6_-SUMO tag was cleaved by incubation with His_6_-Ulp1 protease (2 h, 4°C). Cleaved CHMP7_229-453_ was washed three times with buffer, collected by centrifugation each time, and soluble His_6_-SUMO and His_6_-Ulp1 were discarded in the aqueous supernatant. Cleaved eluate was diluted with buffer to give a final protein concentration of ∼60 µM.

### Analytical Size Exclusion Chromatography

LEM2_1-72_ binding to BAF or BAF-dsDNA complex was assayed by gel filtration chromatography. DNA duplex was prepared as previously described^30^. Combinations of purified LEM2_1-72_, BAF dimers, and DNA duplexes were combined in 1:1:2 molar ratio. Following incubation at room temperature for 30 min, 50 µL of reaction mixtures were applied to a Superose 6 3.2/300 column (GE Life Sciences) in buffer (20 mM HEPES, pH 7.5, 150 mM NaCl, 5 mM DTT, and 10% glycerol) equilibrated at 4°C. The flow rate was 40 µL/min for all experiments. Retention volumes for major peaks absorbing at 280 nm (A280) were recorded. The protein contents of peak fractions were assayed by SDS-PAGE.

### LEM2 Low Complexity Domain Peptides

Chemically synthesized peptides bearing an N-terminal FITC-Ahx modification (Figure S2C) were purchased from from GenScript (Piscataway, NJ), Peptide stock solutions were prepared in milli-Q H2O except LEM2_75-122_ stocks, which were prepared in DMSO. Phase separation of LEM2_75-122_ was induced by dilution in milli-Q H2O or buffer (25 mM HEPES, pH 7.4, 150 mM KCl) to 0.2 mg/mL stock.

### Turbidimetry

Microtubule bundling by LEM2 was quantified by turbidity (absorbance at 340 nm) (Tecan Spark 10M spectrophotometer)^31^. Reactions (10 µL total volume) of purified LEM2 protein or chemically synthesized LEM2 peptides, 1 µM αβ-Tubulin heterodimers polymerized in pre-formed microtubules, KCl (75 mM for LEM2 protein reactions, 0.9 mM for LEM2 peptide reactions), 25 mM HEPES, pH 7.4, 0.5 mM MgCl_2_, 10 µM paclitaxel (Sigma Aldrich), were prepared at room temperature in 384-well non-binding plates (Grendier Bio-One, #781906). Turbidity was measured, with shaking before each read, for reactions in quadruplicate, and averaged for each condition Turbidity values for reactions of LEM2 protein/peptide with microtubules were corrected for the turbidity of LEM2 protein/peptide alone, normalized to turbidity of microtubules alone, and plotted (Turbidity a. u.) against [LEM2]. Sigmoidal curve fitting and half-maximum LEM2 concentrations were calculated with GraphPad Prism software. Error bars are standard error of the mean. Turbidity data was corroborated by complementary negative stain electron microscopy.

### Protein labeling for fluorescence imaging

Proteins were fluorescently labeled for microscopy using Microscale Protein Labeling kits from Thermo Scientific: Alexa Fluor™ 488 (A30006), Alexa Fluor™ 555 (A30007) and Alexa Fluor™ 647 (A30009). LEM2_1-208_ contains a single lysine residue and could not be efficiently labeled with primary amine reactive probes. Instead, His_6_-SUMO-LEM2_1-208_ was labeled with Alexa-488™ for fluorescence microscopy and a minimal amount was used to supplement indicated concentrations of unlabeled LEM2_1-208_ for experiments.

### Phase separation and droplet association assay

LEM2_1-208_ droplet formation was induced at room temperature by reducing the salt concentration to 150 mM KCl, keeping the concentrations of other buffer components the same. Droplets were allowed to form and grow for 15 min before other components were added in tests for droplet association. Droplet association to MT was tested by the addition of 1 µM αβ-Tubulin polymerized into MT, following the Cytoskeleton, Inc. protocol, in a molar ratio of 1:7 HiLyte647-Tubulin:unlabeled Tubulin, in G-PEM buffer (Cytoskeleton Inc. BST01). Unpolymerized tubulin, 1:7 labeled:unlabeled molar ratio, and Alexa555-labeled LEM2_395-503_ were tested for association with LEM2_1-208_ droplets at concentrations of 8 µM each. Note that 150 mM KCl is prohibitive to polymerized tubulin in the absence of stabilizing proteins.

### Spinning Disk Confocal Microscopy with Fluorescence Recovery After Photobleaching (FRAP)

Reactions were prepared in tubes and transferred into a glass-bottom 384-well plate (Grendier Bio-One, #7781892) for imaging. LEM21-208 and LEM275-122 droplets were allowed to settle for 15 min before imaging. Spinning disk confocal microscopy with FRAP was carried out using a W1-CSU with a Borealis upgrade (Andor) and ILE Laser Launch (laser lines 405nm, 488nm, 561nm, 647nm; Andor) on a Nikon Eclipse Ti microscope (Nikon Instruments, Inc.) equipped with a Rapp Optoelectronic UGA-40 (Rapp OptoElectronic) photobleaching unit and controlled though MicroManager 2.0beta (Open Imaging). Fluorescence images were collected with an Andor Zyla 4.2 cMOS Camera (Oxford Instruments). Samples were imaged using a CFI Plan Apochromat VC 100X Oil NA 1.4 objective (Nikon Instruments, Inc.). Excitation wavelengths were: 488 nm for Alex488TM or FITC; 561 nm for Alex555TM; 640 nm for Alex-647. Photobleaching for FRAP experiments was done with a 473 nm laser (Vortran) for 200 ms per bleach event. Images were recorded with a frequency of 1 Hz, starting with at least one image recorded pre-bleach, including the bleach event, and up to 240 s post-bleaching. Partial and whole droplet FRAP experiments were performed in triplicate, and n=17 for the bundled-MT FRAP experiment. The ratio of the intensity within bleached regions of interest to background were calculated with ImageJ for each time point and normalized to the intensity at the time of bleaching. Replicates are aligned to the time of bleaching, averaged, and plotted ± standard deviation.

### Negative stain electron microscopy

Continuous carbon grids (200-400 mesh copper, Quantifoil) were glow-discharged (PELCO EasiGlow, 15 mA, 0.39 mBar, 30 s). Samples (3 – 5 µL) were stained with 0.75% uranyl formate as described previously^25^. Images were collected with a Tecnai T12 microscope (FEI Company, Hillsboro, USA) with a LaB6 filament, operated at 120 kV, and data was captured with a Gatan Ultrascan CCD camera (Gatan, Inc., Pleasanton, USA). For MT-binding assays, reactions were prepared on grids and incubated for 2 min. MT were used at a concentration of 1 µM αβ-Tubulin in pre-formed MT. LEM2_1-208_ was screened for reactivity with MT at concentrations ranging from 0.25 - 4 µM, and LEM2 peptides were screened at concentrations ranging from 10 - 200 µM.

### LEM2-induced CHMP7 Polymerization

LEM2 and CHMP7 proteins were mixed at concentrations ranging from 4 - 8 µM in a total volume of 50 µL. LEM2_395-503_ was present in twofold molar excess to CHMP7 proteins, unless otherwise stated. The buffer was adjusted to between 600 and 800 mM KCl. Reactions were dialyzed for 9-12 h into low-salt buffer (30 mM HEPES, pH 8.0, 25 mM KCl, and 1 mM DTT) using Slide-A-Lyzer MINI Dialysis Device, 10K MWCO (Thermo Fisher Scientific). For experiments including MT, a 3x stock of pre-formed, lyophilized microtubules was prepared in buffer (30 mM HEPES, pH 8.0, 800 mM KCl, 1 mM DTT, 60 uM paclitaxel, and 3 mM MgCl_2_), and dialysis buffer was supplemented with 10 µM paclitaxel and 1 mM MgCl_2_. After dialysis, polymeric assemblies were concentrated by low speed centrifugation (5,000 g, 5 min) and resuspended in 30 µL buffer for negative stain EM, SDS-PAGE, or cross-linking mass spectrometry analysis. To determine the stoichiometry of the CHMP7-LEM2_395-503_ polymer, CHMP7 (4 µM) was titrated with LEM2_395-503_ (0, 0.08, 1 and 4 µM), dialyzed, and polymers wer collected by low-speed centrifugation. The supernatant was collected such that 10 µL where left behind as the pellet fraction. Quantities of polymerized and unpolymerized CHMP7 at each LEM2 condition were measured by Coomassie-stained SDS-PAGE, in which equal volumes of pellet and supernatant fractions were loaded. Intensities were quantified by ImageJ. The data was normalized to samples containing 0 and equimolar amounts of LEM2_395-503_. The experiment was performed in triplicate, the average was calculated, and error bars are standard deviation.

### Liposome preparation

Lipid solutions (Avanti Polar Lipids) were resuspended in chloroform, and liposomes were prepared as previously described^25^. Briefly, lipids (2 mg total) were dried in a glass vial to give final ratio (mole %) of 30% egg phosphatidylserine (PS), 30% egg phosphatidylcholine (PC) and 40% phosphatidylethanolamine (PE). Lipid films were dispersed in buffer (40 mM HEPES, pH 8.0, 150 mM KCl) to produce liposomes at a final concentration of 1 mg/ml, and stored at −80°C.

### Membrane-induced CHMP7 polymerization

CHMP7 was dialyzed or diluted to reduce the salt concentration from 800 mM to 150 mM KCl (supplemented with 5% glycerol, 40 mM HEPES, pH 8.0, 1 mM DTT) to give a final protein concentration of 0.1 mg/ml. Liposomes and CHMP7 were mixed 40:1 (w/w) and negative stain EM grids were prepared immediately.

### Electron Microscopy data acquisition and 2D classification

Membrane-induced CHMP7 polymers were prepared for negative stain EM and imaged with a Tecnai T20 microscope (FEI) with a LaB6 filament, operated at 200 kV. 227 micrographs were collected with a TemCam-F816 8k x 8k camera (TVIPS) using SerialEM software^32^, with a nominal pixel size of 1.57 Å. The defocus was 0.7 −1.7 μm and the total dose was 20 e-/Å^2^. Particles containing a repeating polymeric unit were picked manually along the polymeric protein chain, yielding 6094 particles. Specifically, particles were picked from polymers detached from membrane, which were in a favorable orientation for subsequent classification. Particles were picked and 2D-classified with RELION software.

### Cross-linking Mass spectrometry

Full length CHMP7 (60 µg) was polymerized with equimolar His_6_-SUMO-LEM2_395-503_, following the described polymerization assay, and crosslinked with 2 mM light crosslinker (H_12_-BS3, Creative Molecules) for 30 min at 30°C. His_6_-SUMO-tagged LEM2_395-503_ was used to achieve a higher protein concentration. Full length, monomeric CHMP7 (60 µg) was crosslinked with 2 mM heavy crosslinker (D_12_-BS3, Creative Molecules) under the same conditions. Reactions were quenched (10 mM ammonium bicarbonate, 10 min, room temperature), and light and heavy crosslinked reaction mixtures were combined and processed for mass spectrometry as described previously^33,34^. Crosslinked products were enriched by size-exclusion chromatography (Superdex Peptide, GE Healthcare Life Sciences) as described previously^34^ and fractions eluting between 0.9 and 1.4 mL were dried, resuspended in 0.1% formic acid for MS analysis. The fractions starting at 0.9 ml and 1.3 ml were combined prior to evaporation to make four total SEC fractions.

LC-MS analysis was performed with a Q-Exactive Plus mass spectrometer (Thermo Scientific) coupled with a nanoelectrospray ion source (Easy-Spray, Thermo) and NanoAcquity UPLC system (Waters). Enriched fractions were separated on a 15 cm x 75 μm ID PepMap C18 column (Thermo) using a 60-minute gradient from 5-28% solvent B (A: 0.1% formic acid in water, B: 0.1% formic acid in acetonitrile). Precursor MS scans were measured in the Orbitrap scanning from 350-1500 m/z (mass resolution: 70,000). The ten most intense triply charged or higher precursors were isolated in the quadrupole (isolation window: 4 m/z), dissociated by HCD (normalized collision energy: 24), and the product ion spectra were measured in the Orbitrap (mass resolution: 17,500). A dynamic exclusion window of 20 sec was applied and the automatic gain control targets were set to 3e6 (precursor scan) and 5e4 (product scan).

Peaklists were generated using Proteome Discoverer 2.2 (Thermo), and crosslinked peptides searched for with Protein Prospector 5.20.23^35^. 85 of the most intense peaks from each product ion spectrum were searched against a database containing His_6_-SUMO-LEM2_395-503_ along with the sequences of 10 other proteins comprising CHMP subunits and Tubulin, concatenated with 10 randomized decoy versions of each of these sequences (121 sequences total). Search parameters were: mass tolerance of 20 ppm (precursor) and 30 ppm (product); fixed modifications of carbamidomethylation on cysteine; variable modifications of peptide N-terminal glutamine conversion to pyroglutamate, oxidation of methionine, and “dead-end” modification of lysine and the protein N-terminus by semi-hydrolyzed heavy and light BS3; trypsin specificity was used with 2 missed cleavages and three variable modifications per peptide were allowed. Unique, crosslinked residue pairs were reported at a 1.5% FDR threshold, corresponding to a Score Difference cutoff of 15.

For quantitative analysis, precursor ion filtering in Skyline 4.1^36^ was used to extract light:heavy crosslinked precursor ion signals. Skyline does not directly support crosslinking analysis, so crosslinked peptides were linearized and exported as a spectral library as described in^37^. Transitions were generated for every light or heavy crosslinked peptide species discovered in the Protein Prospector search and paired with its corresponding heavy or light counterpart. Precursor ion transitions matching the first three isotopes were extracted across all four LC-MS fractions. Each extracted ion chromatogram was manually inspected and the start and end points were adjusted to ensure that the correct peaks were detected and that there were no interfering signals. Isotope dot products were required to exceed 0.85. The peak areas were re-imported into R and summarized at the level of crosslinked residues for each light and heavy crosslink. Peak areas were summed for all peptides matching a given crosslink. Finally, log2 ratios of the heavy to light peak areas were determined. Filtered cross-links were mapped to the primary protein structure using xiNET^38^. Quantitative changes are displayed as a cross-link contact map using an in house R-package. The x- y-axes map to protein and residue position, with each dot representing a cross-linked pair. The area of the dot is proportional to the number of spectral counts identifying a cross-link (total number of product ion spectra that match a cross-linked residue pair), while enrichment in either monomeric (heavy BS3 – red) or polymeric (light BS3 – blue) sample is indicated by the color scale.

### Homology modeling and cross-link mapping

Homology models of human CHMP7 and LEM2 domains were created with Phyre2^39^ (Table S1). Reference structures were selected based on confidence scores and homology to reference structure. Models were validated by mapping cross-linking data to the models using UCSF Chimera^40^ together with Xlink analyzer^41^.

### Immunostaining for fluorescence microscopy

HeLa cells were fixed at room temperature in 2% paraformaldehyde for 25 min. The primary antibodies used for immunodetection were rabbit α-IST1^42^, rabbit α-Tubulin (ab18251; abcam), rat α-Tubulin (YL1/2; Accurate Chemical & Scientific), SUN1 (ab124770; abcam), SUN2 (gift from Brian Burke), and mouse α-53BP1 (MAB3802; Millipore). After incubation with fluorescently labeled secondary antibodies (α-rabbit 488, α-mouse 488, α-rabbit 647, and α-rabbit 568; Thermo Fisher), coverslips were mounted with DAPI ProLong Gold (Thermo Fisher) and imaged by either widefield microscopy (Zeiss Axioskop 2; Axiovision software), spinning disk confocal microscopy (Nikon Eclipse T*i*; Metamorph software), or stimulated emission depletion microscopy. For quantifying IST1 localization: α-rabbit 488 against rabbit α-IST1; for assessing DNA Damage: α-mouse 488 against mouse α-53BP1 and α-rabbit 568 against rabbit α-Tubulin; for quantifying the Tubulin and nuclear envelope phenotype: α-mouse 488 against mouse α-SUN2 and α-rabbit 568 against rabbit α-Tubulin; for interphase phenotype: α-rabbit 647 against rabbit α-SUN1 and α-mouse 488 against rat α-Tubulin; for STED: α-mouse 568 against mouse α-Tubulin and α-rabbit 647 against rabbit α-LEM2.

### Stimulated Emission Depletion (STED) microscopy

STED microscopy was performed on a Leica TCS SP8 STED 3X confocal laser-scanning microscope equipped with a HC PL APO CS2 100x/1.40 OIL objective. Confocal sections were imaged with Leica LAS X Core software and processed with Huygens Software Suite (SVI). Images were recorded using 405 nm laser line at 1.4% laser power to image DAPI, and a 572 nm Laser line at 5.6% laser power to image Tubulin in confocal detector mode. LEM2 was imaged with a 653 nm laser line at 2.5% laser power in STED pulsed detector mode (gate start at 0.3 ns and gate end at 6.5 ns) with a Huygens saturation factor of 5.7. Deconvolved Images were further processed in ImageJ (NIH).

### Statistics

For time-lapse colocalization experiments tracking GFP-CHMP7 with either LEM2-mChr or LEM2_Δ415-485_-mChr, data was collected across three independent experiments and individual points corresponding to each cell (two disks) were plotted showing standard deviation (LEM2-mChr: *n*=14; LEM2_Δ415-485_-mChr: *n*=11; see below for method of quantification). For quantification of IST1 recruitment, the mean percent of anaphase disks for each category of IST1 recruitment was determined from three independent experiments and plotted showing standard error of the mean (see below for method of quantification; sample size (*n*) and raw data shown in Supplementary Table 1). For quantification of nuclear accumulation of NLS-3xGFP, data was collected across three independent experiments and the mean plotted showing standard deviation (siControl LEM2-mChr: *n*=18; siLEM2-2 LEM2-mChr: *n*=14; siControl LEM2_Δ415-485_-mChr: *n*=24; siControl LEM2_Δ415-485_-mChr: *n*=6; see below for method of quantification). For quantification of NE-tubulin defects, the mean percent of telophase cells with NE-tubulin defects was determined from three independent experiments and plotted showing standard error of the mean (siControl: *n*=75, 35, 37; siLEM2-1: *n*=72, 35, 37; siLEM2-2: *n*=45, 34, 34). For quantification of DNA damage, the mean percent of telophase cells with ≥5 53BP1 foci was determined from three independent experiments and plotted showing standard error of the mean (siControl: *n*=56, 52, 44; siLEM2-1: *n*=64, 56, 26; siLEM2-2: *n*=74, 50, 40). In all cases, two-tailed unpaired t-test was used to determine p-values, which, unless otherwise specified, are indicated as follows: \**P* < 0.05; \*\**P* < 0.01; \*\*\**P* < 0.001; N.S., not significant.

### Quantification from fluorescence microscopy

For illustration, images of anaphase A and B cells were acquired by widefield microscopy at 100X and adjusted so that background fluorescence in the DNA, IST1, and LEM2-mChr channels were comparable between samples. Raw images acquired by widefield microscopy at 100X were used to score the IST1 phenotype in anaphase A (early) and anaphase B (late). IST1 localization to anaphase chromatin masses was assessed in three independent experiments in which images were randomized and quantified blindly by three independent scorers. Each chromatin disk (two per cell) was scored as having extra robust, robust, weak, or no recruitment for IST1. Robust recruitment was characterized by distinctive foci organized at the core of chromatin masses, whereas weak recruitment was characterized by less intense, often fewer, and less organized foci at the chromatin surface, consistent with what has previously been shown^11^. Extra robust was characterized by strikingly intense IST1 fluorescence, often accompanied by recruitment over most of the chromatin disk. The majority score was used in cases where the three scores differed.

For time-lapse colocalization experiments tracking GFP-CHMP7, images of anaphase cells were acquired by spinning disk confocal microscopy at 60X and were selected for scoring at the time of peak LEM2-mCherry or LEM2_Δ415-485_-mCherry enrichment at the nuclear envelope. In ImageJ, each cell was thresholded for either LEM2-mCherry or LEM2_Δ415-485_-mCherry enrichment. The mean fluorescence (arbitrary units) was measured for each region of interest in the GFP-CHMP7 channel. The regions of interest were subtracted from the area of the whole cell to measure the mean fluorescence of cytoplasmic GFP-CHMP7 for each anaphase cell. The plotted values are the mean GFP-CHMP7 fluorescence of the region of interest, as determined by exogenous LEM2 enrichment, normalized to the cytoplasmic levels of GFP-CHMP7. Plotted points are for individual cells, which were imaged across three different experiments.

Nuclear accumulation of NLS-3xGFP was determined by taking the ratio of nuclear to cytoplasmic NLS-3xGFP at different time points throughout anaphase following imaging by spinning disk confocal microscopy at 60X. In ImageJ, regions of chromatin were defined as regions of interest in the NucBlue ™ channel and used to measure the mean fluorescence (arbitrary units) in the NLS-3xGFP channel. Cytoplasmic levels of NLS-3xGFP were determined by selecting the whole cell, and subsequently deselecting the regions of chromatin. The mean fluorescence of nuclear NLS-3xGFP for each disk was then divided by the mean fluorescence of cytoplasmic NLS-3xGFP of the same cell. Data was collected across three independent experiments and plotted showing standard deviation.

For the purposes of illustration, interphase and telophase cells were acquired by widefield and spinning disk confocal microscopy at 60X and were adjusted so that background fluorescence was comparable between samples. Raw images used to score the nuclear envelope and Tubulin phenotype were acquired by widefield microscopy at 60X in three independent experiments. To score DNA damage, images from three independent experiments were acquired by widefield microscopy at 60X and thresholded uniformly in ImageJ. Nuclear foci were detected using the find maxima function in ImageJ and noise tolerance was held constant for all conditions.

### Time-Lapse Light Microscopy Analysis

For all live imaging experiments images were acquired at 60X and complete cleavage furrow ingression is designated as t=0 minutes.

Stable cell lines used in this study are described in Table S2. Cells stably expressing GFP-Tubulin ^43^ were transiently transfected with siRNA resistant pCMV(Δ5)-LEM2-mCherry (PL13), pCMV(Δ5)-LEM2m21-mCherry (PL14), pCMV(Δ5)-LEM2_Δ43-202_-mCherry (PL15), pCMV(Δ5)-LEM2_Δ145-213_-mCherry (PL16), or pCMV(Δ5)-LEM2_Δ415-485_-mCherry (PL17) using Lipofectamine LTX Plus (Thermo Fisher) for 24 h. Cells were then re-plated on Mat-Tek dishes 8 h before being arrested at G1/S and released, as described below. Twelve hours after release, cells were live-imaged by spinning disk confocal microscopy in the presence of NucBlue™ (Thermo Fisher).

For time-lapse colocalization experiments, HeLa cells stably expressing GFP-CHMP7 incubated overnight. and transiently transfected with either siRNA resistant pCMV(Δ5)-LEM2-mCherry (PL13) or siRNA resistant pCMV(Δ5)-LEM2_Δ415-485_-mCherry (PL17) using Lipofectamine LTX Plus (Thermo Fisher) for 24 h. Cells were then re-plated on Mat-Tek dishes 8 h before being arrested at G1/S and released, as described below. Twelve hours after release, cells were live-imaged by spinning disk confocal microscopy.

To track nuclear integrity, cells stably expressing NLS-3xGFP (PL19) and either siRNA resistant pCMV(Δ5)-LEM2-mCherry (PL13) or pCMV(Δ5)-LEM2_Δ415-485_-mCherry (PL17) were plated on Mat-Tek dishes in the presence of siRNA 8 h before being arrested at G1/S and released, as described below. Twelve hours after release, cells were live-imaged by spinning disk confocal microscopy in the presence of NucBlue™ (Thermo Fisher) in 15 second intervals.

For siRNA depletion in HeLa cells stably expressing H2B-mCherry and GFP-Tubulin, cells were plated on fibronectin coated Mat-Tek dishes in the presence of siRNA 8 h before being arrested at G1/S and released, as described below. Twelve hours after release, cells were live-imaged by spinning disk confocal microscopy.

Cells stably expressing GFP-BAF and LEM2-mCherry were plated on fibronectin coated Mat-Tek dishes 8 h before being arrested at G1/S and released, as described below. Twelve hours after release, cells were live-imaged by spinning disk confocal microscopy.

### siRNA-Mediated Depletion and Cell-Cycle Synchronization

HeLa and U2OS cells were plated on fibronectin-coated coverslips in the presence of 10 nM siRNA oligo, delivered by Lipofectamine RNAiMAX Transfection Reagent (Thermo Fisher). Specific sequences used were: siControl [siScr-1^43,44^], siLEM2-1 [antisense sequence targeting nucleotides 78–98: UUGCGGUAGACAUCCCGGGdTdT^27^], and siLEM2-2 [antisense sequence targeting nucleotides 1,297–1,317: UACAUAUGGAUAGCGCUCCdTdT^27^]. Culture medium containing 2 mM thymidine was then added for 24 h to arrest cells at G1/S. Cells were then rinsed thoroughly with PBS, followed by the addition of culture media. Twelve hours after release, cells were imaged live or fixed for microscopy.

### Immunoblots

To verify efficacy of siRNA treatments and expression of siRNA-resistant constructs, cells were plated in six-well dishes and subjected to the same experimental conditions as those used to generate data. Whole cell lysates were prepared with NP40 lysis buffer, diluted in SDS sample buffer supplemented with DTT, and boiled. Samples were run on 10% SDS-PAGE gels and transferred to PVDF membrane. 3% milk in TBS-T was used for blocking and antibody dilutions. After incubation with primary antibodies [α-LEM2 (HPA017340; Sigma-Aldrich) and α-Tubulin (ab18251; abcam)], reactivity was detected with HRP-coupled secondary antibodies (Thermo Fisher) and chemiluminescence.

---

**Extended Data Table 1.** Template Structures Used for Homology Modeling.

| Protein | AAs covered by model | Template Structure | Model Organism | PDB Identifier |
| --- | --- | --- | --- | --- |
| CHMP7<sub>ESCRTIII</sub> open | 238-356 | SNF7 | *S.c.* | 5FD7 (23) |
| CHMP7<sub>ESCRTIII</sub> closed | 235-373 | CHMP3 | *H.s.* | 2GD5 (51) |
| CHMP7<sub>WH1 + WH2</sub> | 42-223 | VPS25 | *H.S.* | 3CUQ (52) |
| LEM2<sub>WH</sub> | 396-501 | MAN1 | *H.s.* | 2CH0 (20) |

**Extended Data Table 2.**
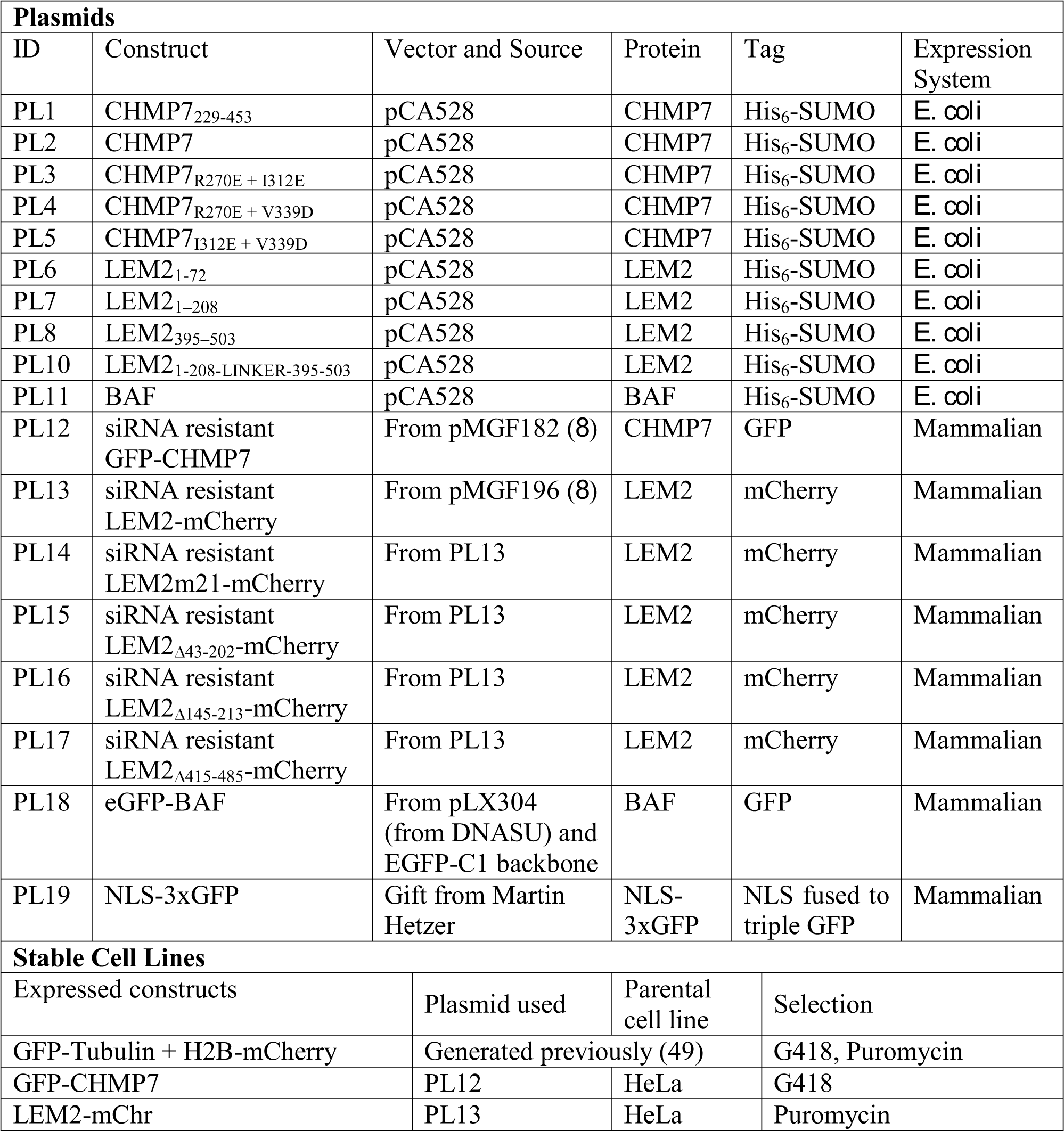
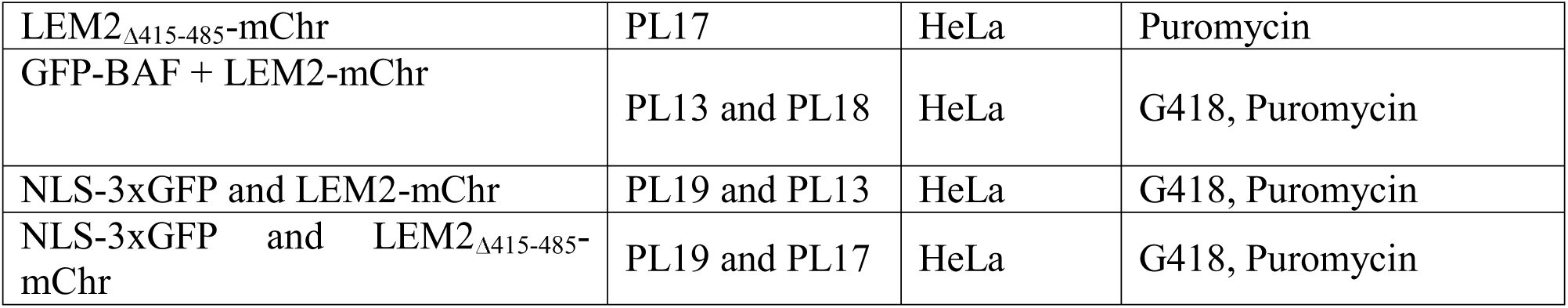
Plasmids and cell lines used in this study. Cell lines stably expressing exogenous, fluorescently labeled proteins were generated by transfecting parental HeLa cells with the described plasmids using Lipofectamine LTX Plus (Thermo Fisher) for 24 h before selection with the specified antibiotics.

| Plasmids |  |  |  |  |  |
| --- | --- | --- | --- | --- | --- |
| ID | Construct | Vector and Source | Protein | Tag | Expression System |
| PL1 | CHMP7<sub>229-453</sub> | pCA528 | CHMP7 | His<sub>6</sub>-SUMO | *E. coli* |
| PL2 | CHMP7 | pCA528 | CHMP7 | His<sub>6</sub>-SUMO | *E. coli* |
| PL3 | CHMP7<sub>R270E + I312E</sub> | pCA528 | CHMP7 | His<sub>6</sub>-SUMO | *E. coli* |
| PL4 | CHMP7<sub>R270E + V339D</sub> | pCA528 | CHMP7 | His<sub>6</sub>-SUMO | *E. coli* |
| PL5 | CHMP7<sub>I312E + V339D</sub> | pCA528 | CHMP7 | His<sub>6</sub>-SUMO | *E. coli* |
| PL6 | LEM2<sub>1-72</sub> | pCA528 | LEM2 | His<sub>6</sub>-SUMO | *E. coli* |
| PL7 | LEM2<sub>1-208</sub> | pCA528 | LEM2 | His<sub>6</sub>-SUMO | *E. coli* |
| PL8 | LEM2<sub>395-503</sub> | pCA528 | LEM2 | His<sub>6</sub>-SUMO | *E. coli* |
| PL10 | LEM2<sub>1-208-LINKER-395-503</sub> | pCA528 | LEM2 | His<sub>6</sub>-SUMO | *E. coli* |
| PL11 | BAF | pCA528 | BAF | His<sub>6</sub>-SUMO | *E. coli* |
| PL12 | siRNA resistant GFP-CHMP7 | From pMGF182 (8) | CHMP7 | GFP | Mammalian |
| PL13 | siRNA resistant LEM2-mCherry | From pMGF196 (8) | LEM2 | mCherry | Mammalian |
| PL14 | siRNA resistant LEM2m21-mCherry | From PL13 | LEM2 | mCherry | Mammalian |
| PL15 | siRNA resistant LEM2<sub>Δ43-202</sub>-mCherry | From PL13 | LEM2 | mCherry | Mammalian |
| PL16 | siRNA resistant LEM2<sub>Δ145-213</sub>-mCherry | From PL13 | LEM2 | mCherry | Mammalian |
| PL17 | siRNA resistant LEM2<sub>Δ415-485</sub>-mCherry | From PL13 | LEM2 | mCherry | Mammalian |
| PL18 | eGFP-BAF | From pLX304 (from DNASU) and EGFP-C1 backbone | BAF | GFP | Mammalian |
| PL19 | NLS-3xGFP | Gift from Martin Hetzer | NLS-3xGFP | NLS fused to triple GFP | Mammalian |
| Stable Cell Lines |  |  |  |  |  |
| Expressed constructs |  | Plasmid used | Parental cell line | Selection |  |
| GFP-Tubulin + H2B-mCherry |  | Generated previously (49) |  | G418, Puromycin |  |
| GFP-CHMP7 |  | PL12 | HeLa | G418 |  |
| LEM2-mChr |  | PL13 | HeLa | Puromycin |  |

|  |  |  |  |
| --- | --- | --- | --- |
| LEM2Δ415-485-mChr | PL17 | HeLa | Puromycin |
| GFP-BAF + LEM2-mChr | PL13 and PL18 | HeLa | G418, Puromycin |
| NLS-3xGFP and LEM2-mChr | PL19 and PL13 | HeLa | G418, Puromycin |
| NLS-3xGFP and LEM2Δ415-485-mChr | PL19 and PL17 | HeLa | G418, Puromycin |

**Extended Data Movie 1. LEM2 droplet formation.**

Droplet formation of 32 uM LEM2_1-208_ was induced by reducing the salt concentration from 500 mM to 150 mM KCl in presence of 0.5 uM Alexa488 labeled His_6_-SUMO-LEM2_1-208._ Sample volume was 40 µl. Images were recorded every min for 120 min total.

**Extended Data Movie 2. Droplet fusion.**

Film clip from ED Movie 1 highlighting

**Extended Data Movie 3. A series of Z-stack images illustrative of the LEM2-depletion phenotype in telophase cells.**

HeLa cells stably expressing H2B-mCherry and GFP-Tubulin were treated with control or LEM2-targeting siRNA and live imaged by spinning disk confocal microscopy.

**Supplementary Table 1. Quantification of IST1 recruitment in mammalian cells.** Numbers reported are the percentage of early and late chromatin discs scored as extra, robust, weak or absent for IST1.

**Supplementary Table 2. Quantified Cross-links from comparative mass spectrometry analysis between monomeric CHMP7 and CHMP7 polymerized by LEM2_395-503_.** Summed peak areas for identified cross-links are listed for Heavy (polymeric CHMP7) and Light (Monomeric CHMP7). In case the identified cross-link was absent in Light sample, the value was set to 40000 for analysis purposes and the log2 ratio is shown as n/a. Cross-links with log2 ratios between 2 and −2 as well as cross-links falling into the region of the His6-SUMO-tag present in LEM2_395-503_ were filtered and not considered in the analysis.

